# The External Microbiome Communicates with the Developing Zebrafish (*Danio rerio*) Embryo Through the Chorion and Influences Developmental Trajectory

**DOI:** 10.1101/2024.05.28.596134

**Authors:** Emily M. Green, Akila Harishchandra, Colin R. Lickwar, John F. Rawls, Richard T. Di Giulio, Nishad Jayasundara

## Abstract

The microbiome has a significant influence on host physiological processes including energy metabolism and neurobiology. However, current knowledge is largely limited to post-embryonic development, highlighting a notable gap in host-microbe communication during embryonic development, particularly in oviparous organisms. This is because the developing embryo is protected from the external environment by the chorion and typically considered to be sterile. We hypothesized the external microbiome influences embryonic development in oviparous organisms despite lack of physical contact with microbes, shaping host physiology beyond embryogenesis. To test this interaction, we utilized zebrafish (*Danio rerio*) reared germ-free or conventionalized with microbes at different times during embryonic development (6 and 24 hours post fertilization) to examine changes in transcriptomics, proteomics, and physiology at 32 hours post-fertilization. In contrast to the prevailing notion, we reveal a significant role of the external aquatic microbial community in regulating embryonic transcript and protein abundance associated with critical developmental processes including energy metabolism and neurodevelopment. Furthermore, we demonstrate the external microbial community drives differential expression of genes involved in cytochrome P450 directed xenobiotic metabolism and associated bioenergetic and behavioral responses following exposure to a CYP1A activator during embryogenesis. These findings reveal embryonic development is an integration of host genetic blueprints and external microbial cues, enhancing knowledge of fundamental developmental processes influenced by embryo-microbe interactions that shape developmental susceptibility to environmental stressors.

**Significance Statement:** Host-microbiome interactions play a crucial role in shaping vertebrate physiology. However, the impact of these interactions during embryonic development remains poorly understood which has limited our evaluation of environmental drivers of developmental disorders and disease. Here, we provide evidence that the external microbiome indirectly communicates with the developing zebrafish (*Danio rerio*) embryo through the chorion, influencing physiological processes including bioenergetics, neurodevelopment, and xenobiotic responses. These findings signify a critical role of the external microbiome during the early stages of embryonic development and may inform research addressing the effects of the maternal microbiome on human embryonic and fetal development, particularly in the context of developmental origins of disease and prenatal chemical exposures.

## 1. Introduction

Animal evolution occurred in an environment abundant with microorganisms, resulting in the emergence of symbiotic relationships that shaped evolutionary trajectory (1). The microbiome, an ecosystem teeming with microorganisms, is critical to maintain normal vertebrate physiology including immune function, digestion, behavior, energy homeostasis, and xenobiotic metabolism (1–11). Furthermore, the microbiome can influence vertebrate development during *post*-embryonic stages (1, 12). However, there is a significant gap in our understanding of host-microbe relationships during embryogenesis, when organisms are highly susceptible to environmental challenges.

Egg-laying oviparous organisms develop surrounded by a diverse microbial environment, although the conventional knowledge is that vertebrate embryos are shielded from physical contact with microbial cells until hatching (9, 13). For example, zebrafish (*Danio rerio)* embryogenesis occurs in a three-layer acellular chorion, perforated with small pore canals spanning 0.5-0.78 μm in diameter. While these pores are essential for oxygen and nutrient exchange, they restrict entry of larger objects such as microbial cells (9, 13–15). Despite the lack of physical contact with microbes, previous studies suggest external bacterial diversity affects hatching and survival rates of oviparous organisms, while providing protection against predation and pathogenic invasion during embryonic development (16–20). Therefore, some vertebrates selectively recruit microbes to the exterior surface of the embryo for protection (9, 21, 22), although the influence of external microbial communities on embryogenesis remain understudied. In mammals, the maternal microbiome influences fetal health, pregnancy outcomes, and pre and postnatal immunity (23– 29). However, the mechanisms by which microbes impact *in utero* development remain poorly understood due to the difficulty in studying these interactions in mammalian models.

Here, using the experimental tractability of zebrafish, we tested the hypothesis that the external microbiome communicates with the developing embryo across the chorion and alters developmental trajectory of the host. Furthermore, given that embryonic development is highly sensitive to external environmental perturbations (21, 30), including toxic chemicals (2, 31), and the well-established role of the microbiome in xenobiotic metabolism of the host (3, 24, 32, 33), we focused on the role of the external microbial community in shaping developmental responses to xenobiotics during embryogenesis. We employed a non-targeted approach and performed RNA sequencing and proteomics on whole embryos at 32 hpf, a critical window of zebrafish development when organ systems, including the nervous system, musculature, vasculature, kidneys, and heart, develop and establish functional roles (34). We created three groups of germ-free (GF) embryos and subsequently conventionalized two of the groups with microbes at 6 hpf (CV6) or 24 hpf (CV24) to investigate molecular pathways influenced by microbial cues from the external environment during embryogenesis. To investigate the link between microbial-driven transcription and translation with physiological consequences of a xenobiotic exposure, we quantified embryonic mitochondrial function and larval behavior. Overall, our findings demonstrate the microbial community is critical for embryonic development and influences developmental responses to xenobiotics.

## 2. Results

### 2.1. The External Microbial Community Drives Transcription in the Developing Embryo

We compared transcriptomic changes between CV6, CV24, and GF embryos to determine the influence of the external microbiome on host gene expression during embryonic development (Figure 1A). All groups were treated to remove the microbial community at 6 hpf and one group maintained under germ-free (GF) conditions. The CV6 group was aqueously conventionalized with a diverse microbial community from recirculating system water immediately following GF derivation at 6 hpf and served as the control. To examine whether the timing of microbial community introduction was important, the CV24 group was created by conventionalizing GF embryos at 24 hpf.

**Figure 1.**
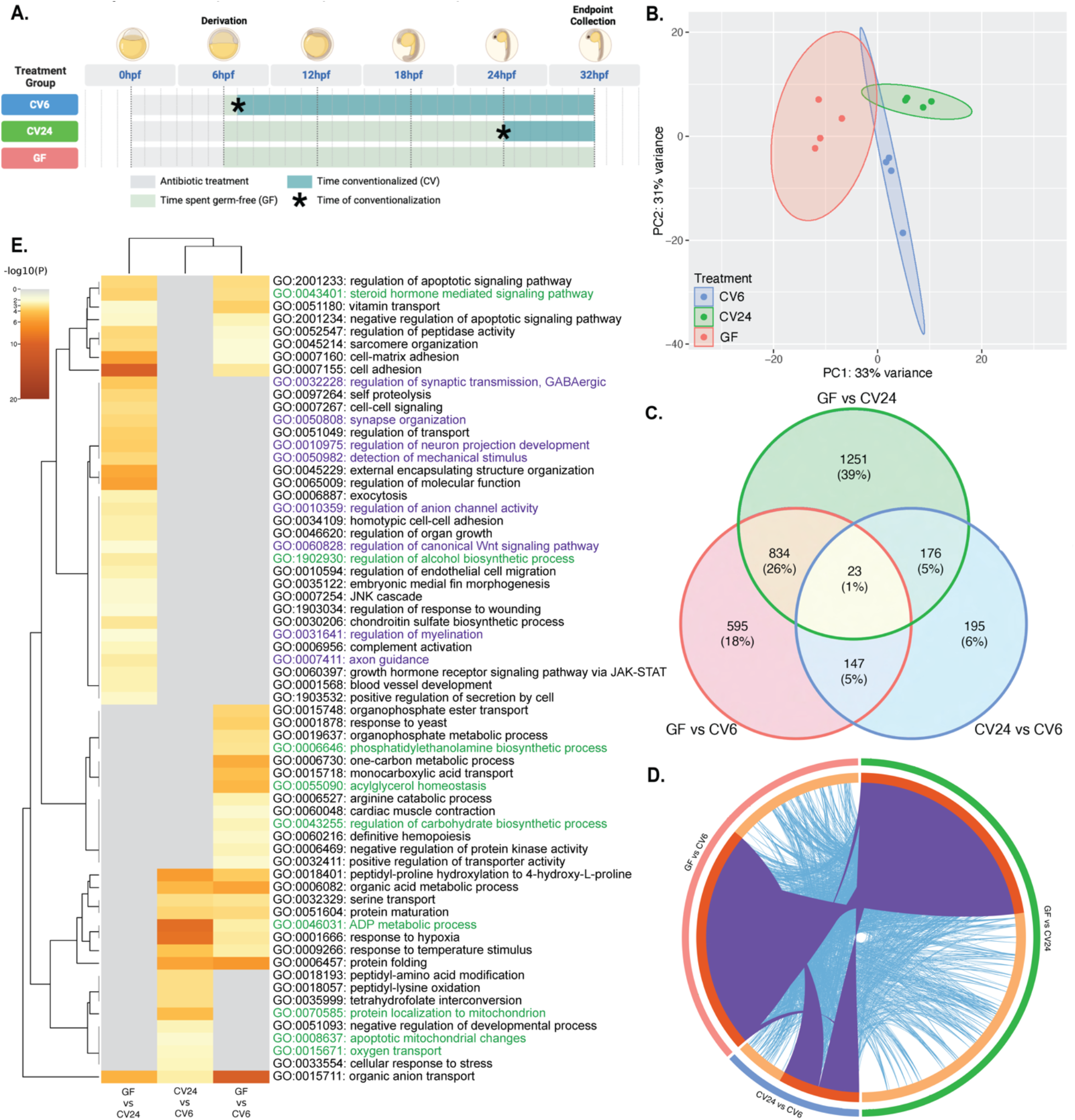
**(A)** Experimental design and treatment groups referenced throughout this study. **(B)** PCA plot of normalized transcript counts for all significant DEGs (adj. p-value<0.05, log_2_Fold Change>|0.5|). Each point represents a sequenced sample (15 embryos) at 32 hpf and ellipses represent ±95% confidence intervals. Treatments are labeled by condition: CV6 (blue), CV24 (green), and GF (pink). **(C)** Venn diagram of significant DEGs across pairwise comparisons from DESeq2 differential expression analysis (adj. p-value<0.05, log_2_Fold Change>|0.5|) (35, 36). The total number of DEGs and the percentage of DEGs compared to the total are shown for each pairwise comparison. **(D)** Circos plot (38) displaying the overlap between gene lists. Purple curves connect identical genes and blue curves connect genes included in the same biological process GO term. **(E)** Heatmap of the most significantly enriched biological process GO terms (y-axis) across pairwise comparisons (x-axis) generated using Metascape analysis (37). Scale represents significance of GO terms (-log_10_p-value). GO terms involved in energy metabolism (lipid metabolism, carbohydrate metabolism, mitochondrial function) and neurodevelopment will be discussed further and are, therefore, colored green and purple, respectively. A full list of all enriched GO terms identified by Metascape across pairwise comparisons can be found in Dataset S2.

At 32 hpf, embryos from each group were randomly selected and used for RNA sequencing (n=4, 15 embryos per sample). Differential gene expression of pairwise comparisons using DESeq2 (35, 36) revealed 3,221 significantly differentially expressed genes (DEGs; adjusted [adj.] p-value<0.05, log_2_Fold Change>|0.5|; Dataset S1). A principal component analysis (PCA) of normalized transcripts illustrated distinct clustering across treatments, demonstrating both microbial colonization status and conventionalization timing affect transcriptional profiles of the developing embryo. Two principal components (PC) were extracted, with PC1 and PC2 describing 33% and 31% of the variance, respectively, representing a combined total of 64% variance (Figure 1B).

Among significant DEGs, 2,284 were differentially enriched in GF compared to CV24 embryos, the greatest number of DEGs across all pairwise comparisons. The GF and CV6 comparison revealed 1,599 DEGs, 857 of which were shared with the GF and CV24 comparison. 541 genes were differentially expressed between CV24 and CV6 embryos, the fewest of any comparison. Of these, 199 were shared with the GF and CV24 comparison and 170 were shared with the GF and CV6 comparison (Figure 1C). Among all pairwise comparisons, the GF and CV6 comparison shared the most DEGs with the GF and CV24 comparison (Figure 1D). Volcano plots revealed the GF vs CV6 and the GF vs CV24 comparisons displayed more DEGs were upregulated in GF embryos (-log_2_Fold Change) compared to CV6 or CV24 embryos. The CV24 vs CV6 comparison revealed significantly more DEGs were upregulated in CV6 embryos (+log_2_Fold Change) compared to CV24 embryos (Fig. S1).

We then used Metascape to probe DEGs from each pairwise comparison and identify enriched biological process gene ontology (GO) terms (Figure 1E) (37). This revealed GO terms involved in lipid metabolism (steroid hormone signaling, alcohol biosynthesis/homeostasis, phospholipid metabolism), carbohydrate metabolism, mitochondrial function (ADP metabolism, protein localization to mitochondrion, mitophagy, oxygen transport), neurodevelopment (GABAergic synaptic transmission, synapse organization, neuron projection development, myelination, axon guidance, anion channel activity, Wnt signaling, detection of mechanical stimulus), immune function (complement activation, growth hormone receptor signaling via JAK-STAT, response to wounding), apoptotic signaling (JNK cascade), protein translation and modifications (peptidase activity, self-proteolysis, arginine catabolism, protein kinase activity, serine transport, protein maturation, protein folding, peptidyl-amino acid modification/oxidation), cell adhesion and signaling (sarcomere organization, endothelial cell migration), and cellular response to stress (organic cyclic compounds, hypoxia, temperature), among others (Figure 1E, Dataset S2).

### 2.2. The External Microbiome Influences Embryonic Transcription of Energy Metabolism and Neurodevelopment Genes

To identify directional trends in expression across treatment groups and interactive effects between conventionalization status and timing of conventionalization, we performed a likelihood ratio test using the DESeq2 LRT function (35, 36), which identified a total of 1,769 significant DEGs (adj. p-value<0.05, log_2_Fold Change>|0.5|) (Fig. S2, Dataset S3). To focus on the processes most significantly impacted within our dataset, these DEGs were subset for further analysis using standard parameters, placing a cutoff at the 1,000 significant LRT DEGs with the greatest differential expression (log_2_Fold Change>|0.64|). This is representative of the trends observed among all significant LRT DEGs (Fig. S3). Four distinct trends in gene expression across treatments were identified. Of those four gene expression patterns, LRT pattern 1 consisted of 306 DEGs downregulated in GF embryos and LRT pattern 2 consisted of 621 DEGs upregulated in GF embryos compared to groups with a microbiome (Figure 2, Dataset S4). Combined, LRT patterns 1 and 2 represented 927 of the top 1,000 LRT DEGs. The remaining patterns contained genes differentially expressed only in CV6 compared to CV24 and GF embryos. LRT pattern 3 was represented by 17 DEGs decreased in CV6 embryos and LRT pattern 4 contained 56 DEGs increased in CV6 embryos, while CV24 and GF embryos displayed comparable expression (Figure 2, Dataset S4). These groups represent 73 LRT DEGs sensitive to the time at which microbial colonization occurred. Notably, no LRT DEGs shared the same trend in GF and CV6 samples and an opposite trend in CV24 samples, confirming expression of all LRT DEGs were driven by the presence of the external microbial community, with a subset of genes sensitive to the timing of microbial cues.

**Figure 2.**
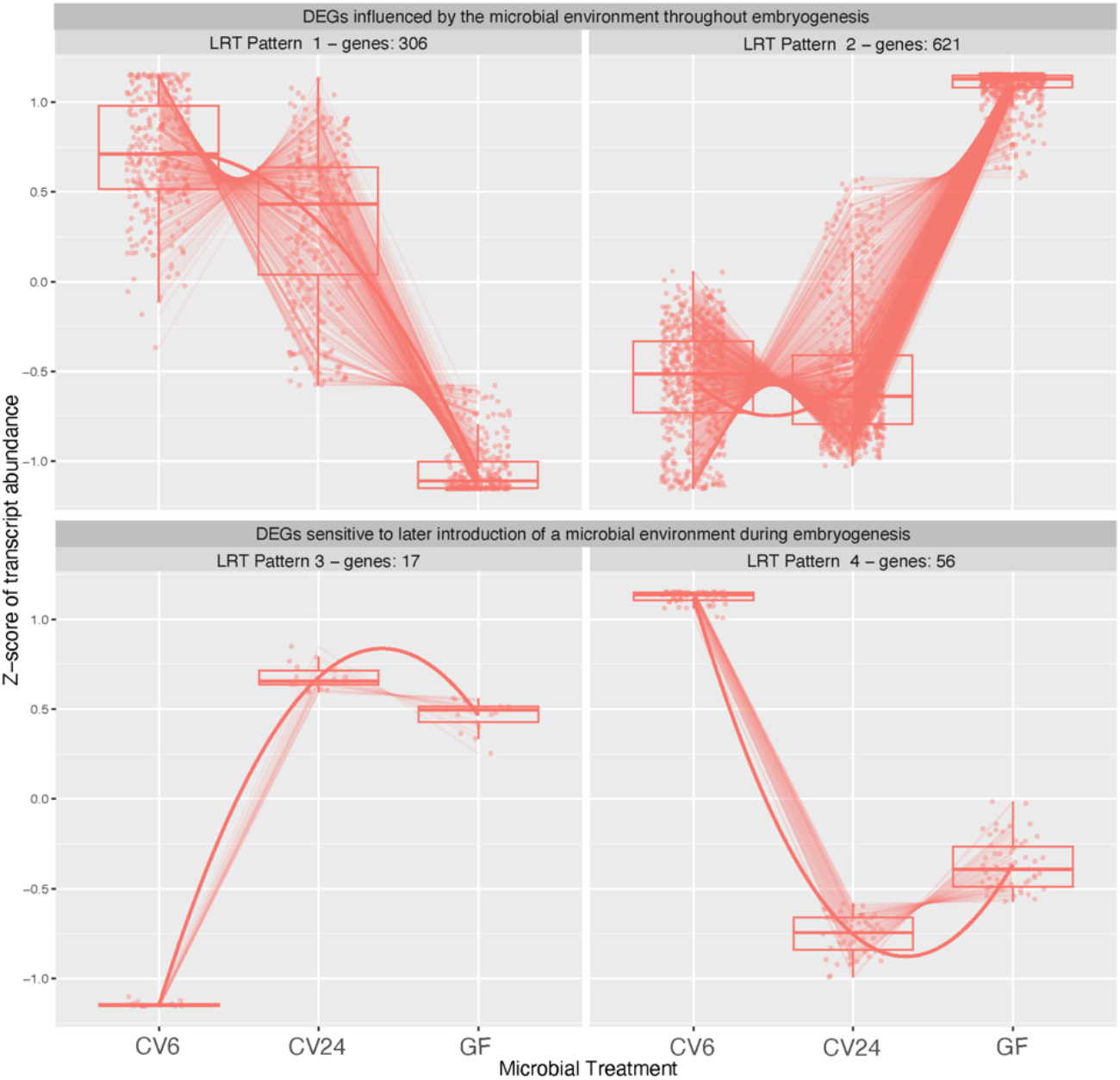
LRT gene expression patterns identified among the top 1,000 DEGs using the DESeq2 LRT function and DEGpatterns (adj. p-value<0.05, log_2_Fold Change>|0.64|) (35, 36, 43). Each pattern represents genes with similar expression across treatments. Dots represent the mean transcript abundance across replicates for a particular gene and transcript counts are scaled by Z-score. Data are displayed by treatment (x-axis) and Z-score of transcript abundance (y-axis).

Since energy metabolism and neurodevelopment were abundant among differentially enriched GO terms across pairwise comparisons, we investigated these processes further. Additionally, energy metabolism and neurodevelopment are frequently assessed as developmental toxicity endpoints, emphasizing the relevance of these processes to our study. All four LRT patterns were analyzed for significantly enriched biological process pathways using the enrichGO function of the ClusterProfiler package (v4.11.0) (39, 40). Among significant pathways (p-value<0.05) involved in processes related to energy metabolism or neurodevelopment, we identified 134 genes involved in energy metabolism (Figure 3A) and 43 genes involved in neurodevelopment (Figure 3B).

**Figure 3.**
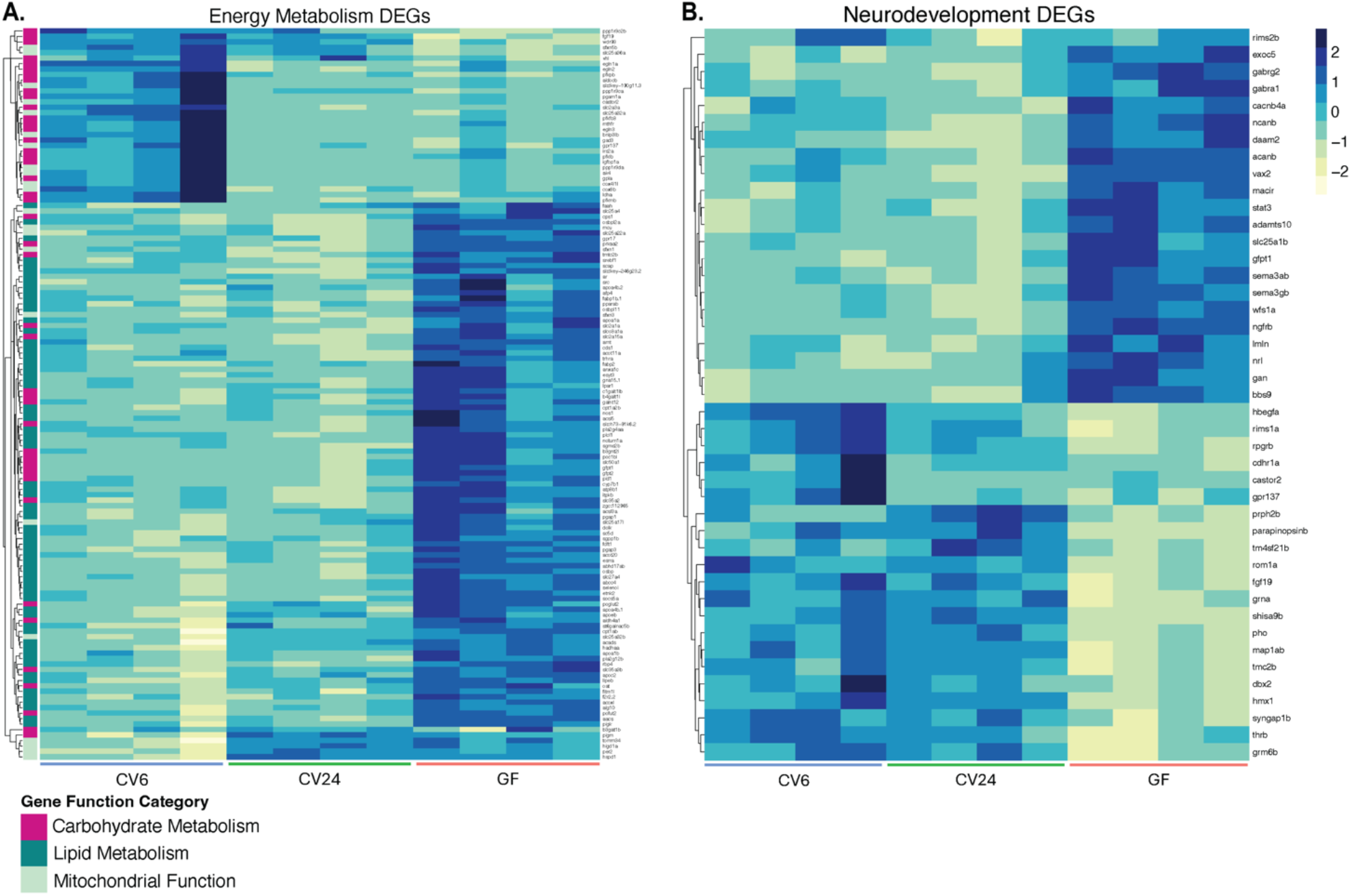
Heatmaps of normalized transcript counts for LRT DEGs involved in **(A)** energy metabolism and **(B)** neurodevelopment. Energy metabolism genes are further categorized by function (carbohydrate metabolism, lipid metabolism, and mitochondrial function) and membership is annotated on the y-axis. Data is displayed by transcript counts for 4 sequenced samples per treatment, scaled by Z-score within rows, (x-axis) and gene name (y-axis). Lists of LRT DEGs included in each heat map are provided in Dataset S6.

Of the 134 energy metabolism genes, 44 were attributed to carbohydrate metabolism, 68 to lipid metabolism, and 22 to mitochondrial function. While these pathways are intrinsically connected, gene function was determined by membership to the assigned significant GO term. Uniquely, all 68 DEGs involved in lipid metabolism were members of LRT Pattern 2 and upregulated in GF embryos compared to embryos with a microbiome (Figure 3A). These genes were involved in lipid localization/transport, lipid homeostasis, fatty acid oxidation, cholesterol metabolism/homeostasis, alcohol metabolism/homeostasis, and steroid metabolism and signaling, among others. Genes involved in carbohydrate metabolism and mitochondrial function were distributed across gene expression trends, suggesting these genes were dependent on both colonization status and the timing of conventionalization. Specifically, genes involved in ADP/ATP metabolism and transport, AMPK activity, and mitochondrial transmembrane transport were upregulated in GF conditions, while genes involved in carbohydrate metabolism, mitochondrial transmembrane transport, and TORC1 signaling were downregulated in GF conditions (Dataset S5). LRT DEGs associated with mitochondrial respiration, mitochondrial organization, mitochondrial membrane permeabilization, ADP/ATP/AMP metabolism, carbohydrate metabolism, protein localization to mitochondrion, and response to reactive oxygen species were differentially expressed in *only* CV6 embryos compared to CV24 and GF embryos (Dataset S5).

Among the 43 genes involved in neurodevelopment, hierarchical clustering revealed approximately half were upregulated and the other half were downregulated in GF embryos, while CV6 and CV24 embryos generally showed comparable transcript counts. This suggested neurodevelopmental gene expression is sensitive to the external microbial environment during embryogenesis, particularly during the second half of embryonic development. We found neurodevelopment LRT DEGs sensitive to microbial colonization to be involved in calcium control of neurotransmitter release and synaptic transmission, retinal/photoreceptor cell differentiation and development, glial cell/neuron differentiation and development, and swimming behavior, among several others (Dataset S5).

Among other processes, we found 33 differentially expressed immune response genes across treatments, a majority of which were upregulated under GF conditions (Fig. S4). Immune function is well-studied in the context of microbial regulation, as gut microbiota produce metabolites such as short-chain fatty acids that regulate inflammatory responses, mobilize immune cells, and defend against microbial pathogens (4, 41, 42). Furthermore, the maternal microbiome influences prenatal and postnatal development of the innate and adaptive immune system in offspring (24, 29). Therefore, the observed enrichment of immune-related genes prior to hatching supports our hypothesis that the external microbial environment impacts development of the embryo. Although many of these genes overlapped with energy metabolism and neurodevelopmental pathways, immune response LRT DEGs were involved in interferon-mediated signaling, eicosanoid transport, oxidative stress, innate immunity, and NF-κB signaling, among others (Fig. S4).

### 2.3. Tissue-Specific Enrichment of Microbial-Driven DEGs

To determine tissue-specificity of overall gene expression in the developing embryo across treatments, we compared our dataset to 31 tissue-enriched gene lists from a published embryonic single-cell transcriptomic dataset (44) as similarly described (45). We utilized the mroast function of the Limma package (v3.60.3) (46) to identify differential expression within each tissue-enriched gene list and assigned each a rank based on significance (FDR adj. p-value<0.05). Each pairwise comparison was represented by DEGs from all 31 tissue-enriched gene lists. The GF and CV6 comparison was most significantly represented by genes enriched in liver, myeloid lineage, integument, intestine, and muscle tissue development, all of which were upregulated in GF embryos compared to CV6 embryos (Figure 4). The GF and CV24 comparison had the most significant representation from gene sets enriched in pharyngeal endoderm, lens placode, intestine, integument, and lateral plate mesoderm, all represented by a downregulation in GF compared to CV24 embryos (Figure 4). Furthermore, differential expression among the CV24 and CV6 comparison was most significantly enriched in the central nervous system, placode, pharyngeal endoderm, germ cell, and intestine, all of which were upregulated in CV24 compared to CV6 embryos (Figure 4). This suggests the influence of the external microbial environment on differential gene expression during embryogenesis targets a broad range of tissue types critical to healthy embryonic development.

**Figure 4.**
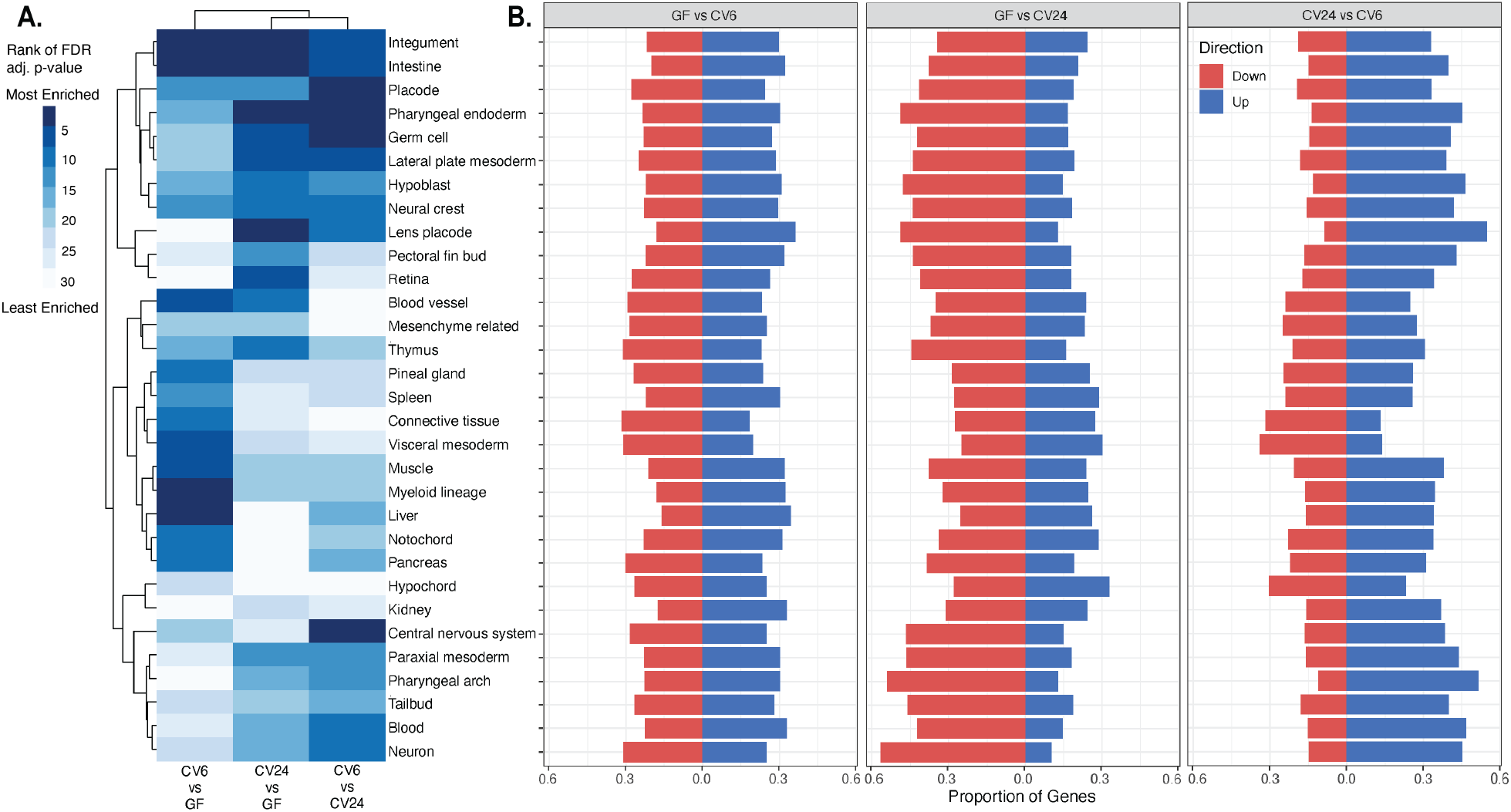
Tissue-enrichment of differentially expressed genes identified using the mroast function from the Limma package (v3.60.3) and a single-cell transcriptomic dataset (44, 46). **(A)** Heatmap of significance of differential expression among tissue-enriched gene sets. Pairwise comparisons (x-axis) and tissue type (y-axis) are shown. Scale represents the rank of all 31 tissue-enriched gene sets determined by the FDR adj. p-value for each pairwise comparison. **(B)** Bar charts displaying the proportion of DEGs downregulated and upregulated (z-score>|1.41|) within each tissue-enriched gene set. Pairwise comparisons are shown in separate panels and data is displayed by the proportion of genes (x-axis) and tissue type (y-axis). Differential regulation is further defined by downregulation (red) and upregulation (blue).

### 2.4. The External Microbiome Impacts Embryonic Abundance of Energy Metabolism and Neurodevelopment Proteins

To determine whether the external microbial environment elicited embryonic proteome changes, an additional subset of CV6, CV24, and GF embryos were randomly selected for protein quantification by LC/MSMS (n=4, 15 embryos per sample). Differential protein analysis using DEP (47) revealed 1,146 differentially abundant proteins (DAPs; p-value<0.05, logFold Change>|0.5|) across all pairwise comparisons (Dataset S7). Of these 492 differed between GF and CV6 embryos and 469 differed between GF and CV24 embryos, 128 of which overlapped with the GF and CV6 comparison. We identified 575 DAPs between CV24 and CV6 embryos, the greatest of all comparisons, 138 of which overlapped with the GF and CV6 comparison and 128 of which overlapped with the GF and CV24 comparison (Figure 5). A PCA revealed 21.3% of variance in PC3 and PC4 is largely explained by microbial colonization status (Fig. S5).

**Figure 5.**
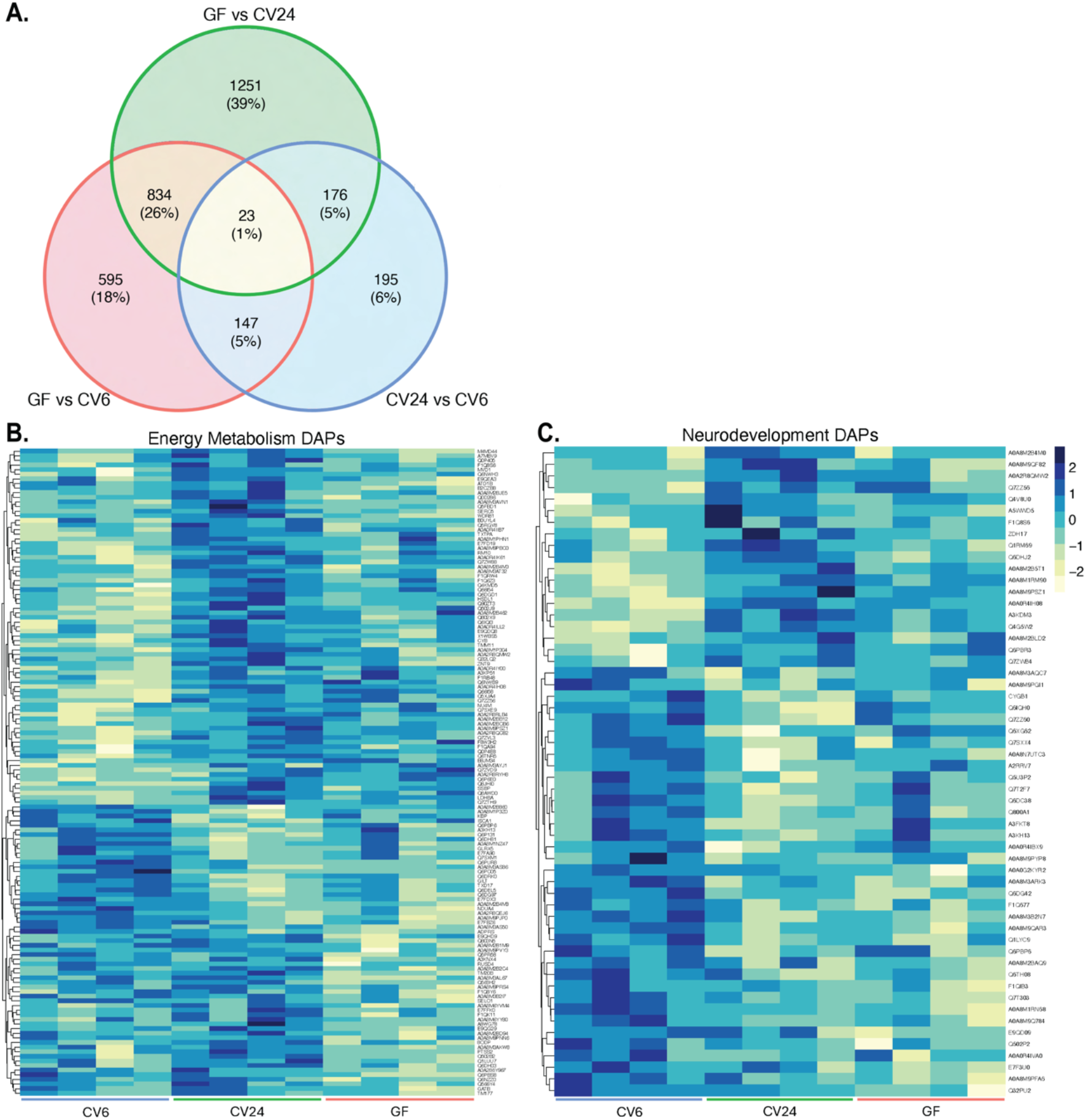
**(A)** Venn diagram of DAPs from DEP pairwise comparisons (p-value<0.05, logFold Change>|.5|). The total number of DAPs and the percentage of DAPs compared to the total are shown for each pairwise comparison. **(B)** Heatmap of protein abundance for DAPs involved in energy metabolism and **(C)** heatmap of protein abundance for DAPs involved in neurodevelopment. Data is displayed by protein abundance for 4 sequenced samples per treatment, scaled by Z-score within rows, (x-axis) and lists of DAPs included in each heat map are provided in Dataset S8.

The DAVID Functional Annotation Tool (48, 49) was used to screen pairwise lists of DAPs for significantly enriched pathways and revealed 140 DAPs involved in energy metabolism (Figure 5B) and 56 DAPs involved in neurodevelopment (Figure 5C). Specifically, the GF and CV6 comparison revealed enrichment of proteins involved in the mitochondrial inner membrane, the TCA cycle, Acyl CoA, ATP synthesis, and respiratory electron transport, while the GF and CV24 comparison revealed differences in the mitochondrial inner membrane, ATPase activity, mitochondrial translation, and phospholipid metabolism (Figure 5B, Dataset S8). Comparing CV24 and CV6 embryos additionally revealed differential abundance of proteins involved in the mitochondrial inner and outer membrane, membrane translocation and protein import, mitochondrial ribosomal subunits and translation, redox cycling, ATP biosynthesis, mitochondrial electron transport chain complex I, oxidative phosphorylation, mTOR signaling, and lipid metabolism (Figure 5B, Dataset S8). Many of the DAPs identified here are involved in similar biological functions to the LRT DEGs identified from the transcriptional dataset, highlighting energy metabolism as a critical embryonic developmental process influenced by the external microbial environment (Figure 5B, Dataset S8). Notably, our results suggest an interaction between cytochrome c oxidase and the microbial environment at gene and protein levels in GF and CV24 embryos (Fig. S6).

The 56 DAPs involved in significantly enriched neurodevelopmental pathways also shared similar functions to the identified LRT DEGs from our transcriptional dataset. Proteins with differential abundance between CV6 and GF embryos were involved in postsynaptic neurons and axonogenesis, while the CV24 and GF pairwise comparison revealed proteins involved in neural crest cell differentiation and calcium signaling (Figure 5C, Dataset S8). The CV24 and CV6 embryos differed in proteins involved in axonogenesis, calcium signaling, and neural crest cell development (Figure 5C, Dataset S8), further highlighting the critical importance of the microbiome during the first half embryonic development in calcium signaling and proper development of axons and neural crest cells.

### 2.5. The External Microbiome Impacts Bioenergetic and Behavioral Responses to an Embryonic Xenobiotic Exposure

Cytochrome P450s are a conserved class of enzymes that play a vital role in the oxidative metabolism of xenobiotics via the aryl hydrocarbon receptor (AhR) (50). As mentioned, our transcriptomic results revealed treatment-specific differential expression of genes involved in cellular responses to cyclic organic compounds, including several xenobiotic metabolism genes (Fig. S7A), with cytochrome P450 1A (*cyp1a*) being the single most upregulated DEG in all treatments with a microbiome (Figure 6A), which was validated via qPCR (Fig. S8). CYP1A protein abundance was also differentially regulated across treatments (Figure 6B), although we observed a discordance in protein and gene expression data. While AhR was not differentially expressed, GF embryos displayed increased aryl hydrocarbon receptor nuclear translocator (*arnt*) expression (Fig. S7A), which is required for translocation of the AhR to the nucleus to induce transcription of downstream genes, such as *cyp1a*. This may be explained as a compensatory mechanism in response to the reduced basal expression of *cyp1a* among GF embryos.

**Figure 6.**
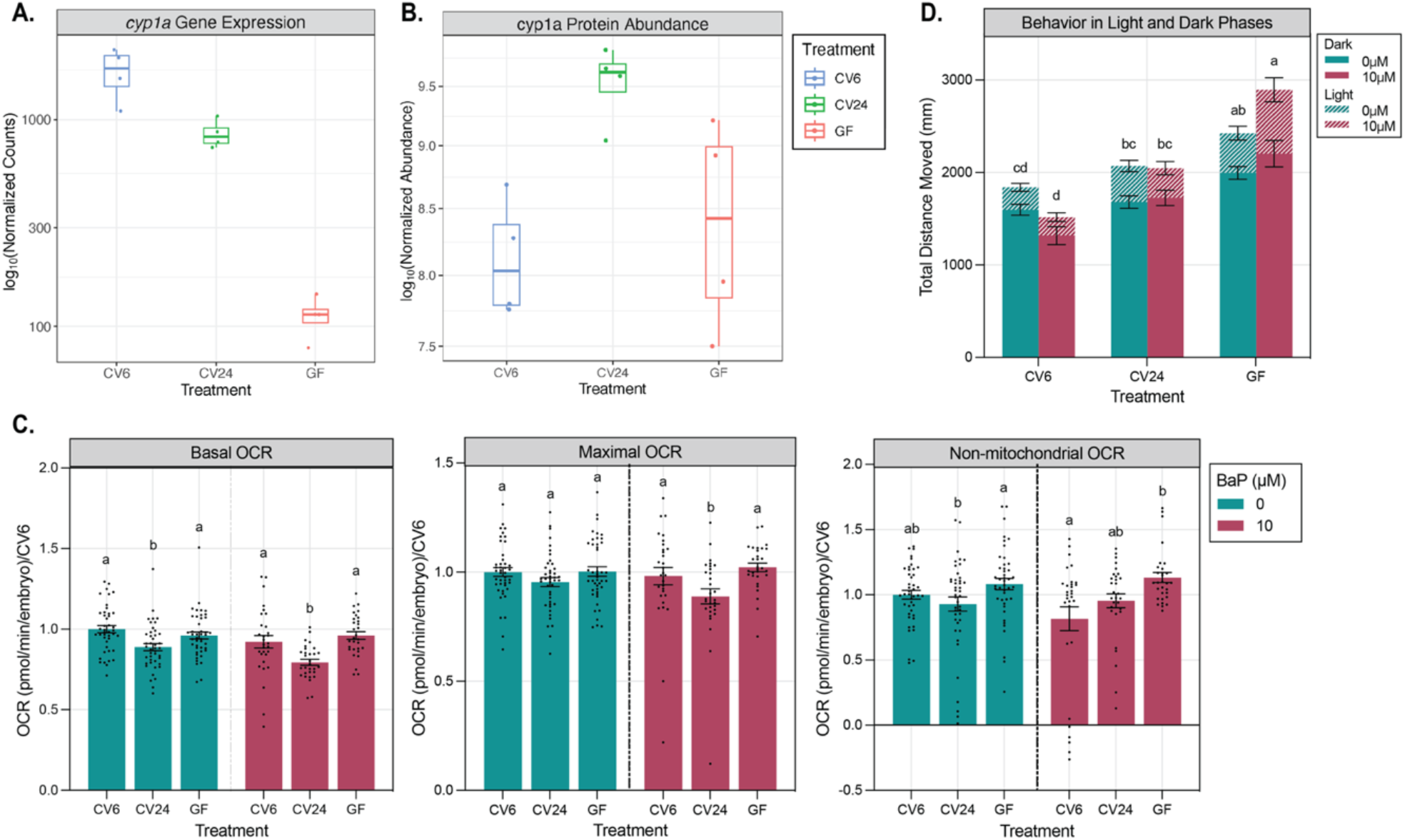
Impacts of the external microbiome on bioenergetic and behavioral responses to an embryonic xenobiotic exposure. **(A, B)** *cyp1a* gene expression **(A)** and CYP1a protein abundance **(B)** in CV6, CV24, and GF embryos at 32hpf. Each point represents a sample containing 15 pooled embryos and are labeled by treatment: CV6 (blue), CV24 (green), and GF (pink). Data are displayed by microbial treatment (x-axis) and log_10_(Normalized Counts/Abundance) (y-axis). **(C)** Basal, maximal, and non-mitochondrial oxygen consumption rates (OCR) in CV6, CV24, and GF embryos at 32 hpf exposed to a vehicle control (Blue, 0.1% DMSO) or 10 μM BaP (red). Significance is represented by different letters and determined by two 1-way ANOVAs, separated by BaP treatment, Tukey’s HSD, p<0.05. Error bars represent standard error of the mean. n=30-44. Data are displayed by microbial treatment (x-axis) and OCR (pmol/min/embryo), normalized to the average of CV6 (y-axis). **(D)** Total distance moved over two sets of alternating 10-minute light/dark cycles in CV6, CV24, and GF larvae at 5 dpf dosed with either a vehicle control (0.1% DMSO) or 10 μM BaP. Significance is represented by different letters and determined by a two-way ANOVA, Tukey’s HSD, p<0.05. Error bars represent standard error of the mean. n=35-64. Data are displayed by microbial treatment (x-axis) and total distance moved (mm) (y-axis). Each condition is labeled by BaP treatment (0 μM – blue, 10 μM – red) and distance moved in light or dark phases (Dark – solid bar, Light – Striped bar).

To assess the developmental requirement of *cyp1a* activation by microbes, we included two additional time points of conventionalization, 12 hpf (CV12) and 30 hpf (CV30) and compared *cyp1a* expression to CV6, CV24, and GF embryos at 32 hpf using qPCR. The CV30 group was conventionalized with microbes two hours prior to endpoint collection to test how rapidly *cyp1a* expression in the developing embryo changed in response to the external microbial environment. Our results confirmed *cyp1a* expression decreased in GF embryos and increased in the presence of a microbial community, regardless of the timing of conventionalization (Fig. S8).

In addition to AhR-directed xenobiotic metabolism, the pregnane X receptor (PXR) is well-known for sensing and metabolizing xenobiotics (51). While activation of the AhR induces genes involved in phase I metabolism, activation of the PXR stimulates phase I and II metabolism genes (51). PXR (*nr1i2*) was significantly enriched in the GF and CV24 pairwise comparison, although differential expression between CV6 and GF embryos was just outside our significance cutoff (log_2_Fold Change=|0.39|), our results suggest expression of *nr1i2* increases in GF embryos during early development (Fig. S7B). This proposes alternative xenobiotic metabolism pathways GF embryos may utilize to compensate for reduced expression of *cyp1a* in the absence of microbial cues.

To determine whether microbial driven differences in *cyp1a* expression influenced physiological responses to xenobiotic exposure, we assessed the effect of the embryonic microbiome on bioenergetic and behavioral responses following exposure to a well-studied AhR agonist and CYP1A inducer, benzo(a)pyrene (BaP) (52, 53). At 32 hpf, *cyp1a* expression is present at basal levels in the central nervous system, endoderm, hematopoietic system, mesoderm, mesenchyme, neural crest, notochord, and periderm (54) and increased *cyp1a* expression is a biomarker of BaP exposure (55). Furthermore, BaP elicits toxic effects on neurodevelopment and mitochondrial function (56). Therefore, at 24 hpf, we exposed CV6, CV24, and GF embryos to 10 μM of BaP or a vehicle control (0.1% dimethyl sulfoxide [DMSO]) for the duration of the experiment. Since transcriptomic results suggested energy metabolism and neurodevelopment were altered across treatments, we assessed embryonic mitochondrial respiration at 32 hpf and larval swimming behavior at 5 days post-fertilization (dpf).

Our results illustrated basal oxygen consumption rates (OCR) decreased in CV24 embryos compared to CV6 and GF embryos, while no changes in maximal OCR were observed in the absence of a BaP exposure (Figure 6C). However, both basal and maximal respiration decreased in CV24 embryos in the presence of 10 μM BaP exposure (Figure 6C), while spare capacity (Fig. S9) and total mitochondrial abundance remained unchanged (Fig. S10). Non-mitochondrial OCR significantly increased in GF embryos compared to CV24 embryos in the absence of a BaP exposure and increased in GF embryos compared to CV6 embryos in the presence of BaP (Figure 6C).

To assess whether the external embryonic microbiome differentially influenced larval behavioral response to BaP exposure, we measured total distance moved during alternating light and dark cycles at 5 dpf. Larvae raised under GF conditions increased total distance traveled (hyperactivity) compared to CV6 larvae. However, CV24 larvae were not statistically different from CV6 or GF larvae and exhibited minor, hyperactivity compared to CV6 larvae (Figure 6D, Fig. S11A). With embryonic 10 μM BaP exposure, total distance moved in both the CV24 *and* GF groups significantly increased compared to CV6 larvae, suggesting the timing of conventionalization is critical for the behavioral response to BaP (Figure 6D). Notably, GF larvae experienced hyperactivity in both the light and dark phases, while CV24 larvae only exhibited hyperactivity in the dark in response to a 10 μM BaP exposure (Fig. S11B).

## 3. Discussion

### 3.1. The External Microbial Environment Affects Embryonic Development

Contrary to the notion that zebrafish embryos are shielded from the external microbial environment (9, 13), our transcriptomic and proteomic data show a distinct interaction between the external microbial environment and embryogenesis. Gnotobiotic studies commonly conventionalize zebrafish around 72 hpf, after hatching (13). However, our study emphasizes the timing of conventionalization is an important consideration when designing future gnotobiotic studies, as the embryo undergoes critical developmental processes sensitive to microbial cues from the external microbial environment. We show the presence of a microbiome is critical for optimal transcriptomic patterning during embryonic development. Furthermore, we identify a small subset of genes (∼7% as shown in the LTR analysis) more similar between CV24 and GF embryos than to embryos colonized earlier in development (CV6), indicating the importance of timing of microbial cues from the external environment. Further studies are needed to determine the drivers of these gene expression patterns and whether gene expression is more dependent on the developmental timing of microbial introduction or the temporal duration of exposure to a microbial community. Further studies are also necessary to examine discrepancies between specific changes in gene and protein abundances, which may be attributed to differences in the speed at which transcription and translation respond to microbial cues.

### 3.2. The External Microbiome Impacts Embryonic Energy Metabolism

It is well established that the gut microbiome influences metabolic homeostasis during larval development in zebrafish, although our findings are among the first to show microbial-directed effects on energy metabolism begin during embryogenesis. Microbial metabolites can serve as an energy source, act as epigenetic modifiers, and bind to G-protein coupled receptors on epithelial cells, affecting lipid and glucose metabolism as well as mitochondrial function during larval development (41, 57–61). Therefore, it is possible these metabolites may play a similar role in influencing energy metabolism during embryonic development.

The overall influence of the external microbiome on embryonic energy metabolism may be attributed to mitochondrial dysfunction. Previous studies during post-embryonic development show microbial metabolites can influence mitochondrial respiration, biogenesis, oxidative phosphorylation, the TCA cycle, ATP production, proton leak, and AMPK activity (59, 62–65). Our results showed treatment specific differences in embryonic expression of genes and proteins involved in mitochondrial function (e.g., TCA cycle, ATP synthesis). Our transcriptomic results further revealed an increase in AMPK (*prkaa2*) in GF embryos compared to embryos with a microbiome, potentially suggesting GF embryos may enhance glucose uptake, mitochondrial fatty acid oxidation, and respiratory capacity through AMPK-dependent mechanisms (62, 66). While mitochondrial content remained consistent across treatments, we observed an increase in *bnip3lb*, a gene involved in mitophagy regulation, in embryos colonized earliest in development. Induction of mitophagy may contribute to maintaining improved mitochondrial efficiency in the presence of a microbial environment during early embryogenesis (67, 68).

In concordance with gene and protein expression, we observed a decrease in basal mitochondrial respiration in embryos that did not receive a microbiome until halfway through embryonic development (CV24). However, it is unclear why mitochondrial respiration was not affected in GF embryos. While GF embryos increased non-mitochondrial respiration rates, it is possible GF embryos reach metabolic homeostasis or shifted energy substrates, possibly aided by AMPK upregulation, in the absence of microbial-directed cues. In contrast, CV24 embryos received microbial-directed cues, although developmentally later than CV6 embryos, and may have adjusted mitochondrial respiration efficiency following those cues.

The observed uniform upregulation of 68 LRT DEGs involved in lipid metabolism in GF embryos may possibly be a compensatory attempt to increase energy metabolism in the absence of microbial-directed cues. Our results demonstrate GF embryos increased expression of genes involved in fatty acid synthesis and oxidation, cholesterol synthesis, and lipid absorption/transport, notably *nr1i2* (PXR) and *apoa1*. Induction of *apoa1* via PXR is linked to lipid accumulation, and beta-oxidation of lipids (51, 69, 70) and may explain the observed differences in lipid metabolism genes. Germ-free studies during post-embryonic development suggest cholesterol synthesis and fatty acid oxidation increase under GF conditions (41, 61, 71, 72), while fatty acid synthesis, absorption, transport, and accumulation decrease under GF conditions (61, 62, 71–73). It is possible in the absence of a microbiome during early development, lipid synthesis is initially upregulated, as we see at 32 hpf, and subsequently downregulated as energy storage becomes depleted. In summary, our data clearly shows the external microbiome may influence energy homeostasis during embryogenesis via multiple mechanisms or through one master regulator. Further studies are needed to investigate the mechanisms by which energy dynamics shift in the absence of microbial-directed cues during embryogenesis.

### 3.3. The External Microbiome Impacts Embryonic Neurodevelopment

The bidirectional communication of the gut-brain axis is well characterized during post-embryonic development as microbes and their products transmit signals from the gut to the brain (5, 10, 71, 74–77). While calcium and neurotransmitter signaling, synaptic plasticity, visual development, axonogenesis, and HPA/HPI axis activation were previously investigated in the context of microbial regulation during *post*-embryonic development (5, 7, 26, 78–80), microbial regulation of these pathways during embryonic development is less understood. Our findings support a model in which the external microbial environment influences these critical neurodevelopment pathways prior to hatching. Utilizing a mouse model, Vuong et al. (2020) demonstrated the maternal microbiome modulates fetal neurodevelopment, resulting in post-natal behavioral effects. This study discovered axonogenesis was reduced during embryogenesis in offspring from microbiome depleted dams, marked by differential expression of GABA receptor genes *gabra1* and *gabrg2* (26). Similarly, our study observed differential regulation of genes and proteins involved in axonogenesis, including *gabra1* and *gabrg2* and p21 protein (Cdc42/Rac)-activated kinase 1 and 2a (PAK1 and PAK2a). These studies collectively demonstrate the role of the microbiome in development commences prenatally during embryogenesis, a developmental time point largely overlooked in microbiome studies. We also demonstrated genes associated with the central nervous system were the most significantly differentially expressed among the CV24 and CV6 pairwise comparison, further emphasizing an important role in microbial-directed neurodevelopment during the first 24 hours of embryogenesis.

Findings from Rea et al. (2022) also aligned with our results, suggesting a delay in zebrafish neurodevelopment occurs under GF conditions and is rescued by the addition of specific microbes or their metabolites. Rea et al. (2022) suggests GF embryos experience dysregulated eye and lens development, Wnt signaling, and neuron and glial organization. Our results confirm disruption of eye and lens development and neurogenesis, although the specific DEGs involved in these processes differed, which may be attributed to differences in collection timing or methods employed in analysis. Our studies were conducted at 32 hpf, while Rea et al. (2022) investigated differences at 48 hpf. Some DEGs were noted to not appear until after hatching, suggesting timing of collection may play a role (80). While dysregulated Wnt signaling pathways were not significant in our LRT analysis, Wnt signaling was identified as one of the most significantly enriched GO terms among the GF and CV24 pairwise comparison.

Our larval behavioral studies complement the embryonic transcriptional and translational differences. While larval behavioral changes are in concordance with previous studies (5, 7, 9, 32), we additionally demonstrated a time-dependent impact of conventionalization, as CV24 display mild hyperactivity, but not to the extent of GF embryos. Collectively, these studies support a role of the external microbial community in modulating neurodevelopment during embryogenesis, although further research is needed to elucidate the mechanisms.

### 3.4. The External Microbiome Impacts Embryonic Responses to Xenobiotic Exposure

Despite lack of fully functioning livers until 4 dpf (81), zebrafish embryos are capable of xenobiotic metabolism during embryogenesis (82). We proposed a possible mechanism by which the microbiome influences embryonic responses to chemical exposure, supported by differential regulation of genes involved in xenobiotic metabolism, including several CYPs, across treatments. While previous studies show microbial metabolites activate AhR, inducing transcription of *cyp1a* (83–86), we observed *cyp1a* was the most upregulated gene in the presence of a microbiome, further suggesting microbial metabolites may elicit impacts on the developing embryo. However, the discordance in *cyp1a* gene expression with protein abundances warrant further studies and may be attributed to differences in post-translational modifications of CYP1A across treatments. CYP1A enzyme activity can be observed as early as 8 hpf (87) and cytochrome P450 transcripts and proteins follow temporally dynamic expression patterns to during embryogenesis to direct normal developmental processes (82, 87, 88). Therefore, it is possible microbial cues may offset normal fluctuations.

We observed *nr1i2* (PXR) increased in the absence of a microbial community during early development. Furthermore, a corresponding increase of phase II metabolism genes, including glutathione S-transferases (*gstt2, gsta.2*) and a decrease in sulfotransferases (*sult6b1*) were observed in GF embryos, suggesting a proposed mechanism by which the external microbial environment modulates xenobiotic metabolism in the developing embryo. Similar to the AhR, xenobiotics, dietary compounds, and microbial metabolites, such as indoles, are ligands of PXR (89). Therefore, it is unclear how PXR is transcriptionally enriched and future studies may investigate the mechanisms by which PXR is induced in a germ-free environment.

Our results also demonstrate the external microbial environment impacts host physiology in response to an embryonic xenobiotic exposure. While our results showed basal mitochondrial respiration decreased in embryos lacking a microbiome until halfway through embryonic development (CV24), this difference became pronounced in the presence of a xenobiotic exposure. With exposure to BaP, CV24 embryos showed an overall reduction of both basal *and* maximal mitochondrial respiration compared to CV6 and GF embryos. BaP is previously reported to affect post-embryonic zebrafish bioenergetics (56). However, our study is the first to reveal the microbiome is increasingly important during the first 24 hours of development to maintain mitochondrial function when challenged with a xenobiotic exposure.

In terms of swimming behavior, while only GF larvae experienced significant hyperactivity in the absence of the BaP exposure, a time-dependent impact of conventionalization became evident upon exposure to BaP. When exposed to BaP, both GF *and* CV24 larvae displayed hyperactivity compared to CV6 embryos conventionalized earlier during development. While BaP is known to impact larval behavior (56), our study emphasized the significance of the external microbial community during the first 24 hours of development in regulating behavioral responses to BaP exposure. While previous studies illustrated microbiota impacts neurodevelopmental toxicity of xenobiotics during larval development (32), we further demonstrated the presence of an external microbiome during embryogenesis and the *timing* of introduction was significant for larval behavioral responses to xenobiotics, suggesting the external microbial environment directly influenced neurodevelopment during embryogenesis.

## 4. Conclusion

The current data clearly demonstrate host-microbiome interactions commence as early as embryogenesis, shaping the trajectory of organismal development. The presence of the external microbial community and the timing of conventionalization impacts key processes including embryonic energy metabolism, neurodevelopment, and responses to a xenobiotic exposure. This transforms our approach to germ-free studies, particularly in the context of embryonic development, and challenges the current notion that oviparous embryos develop completely shielded from the microbial environment until after hatching. Current findings are also of particular significance to the developmental plasticity and fitness of oviparous organisms in microbially dynamic natural habitats. Our findings expand understanding of host-microbe interactions during the earliest stages of vertebrate development. This will provide a foundation to examine the role of the external microbiome in developmental origins of disease as well as the developmental effects of *in utero* and *in ovo* chemical exposures.

## 5. Methods

### 5.1. Zebrafish husbandry

Adult Ekkwill zebrafish (Ekkwill Waterlife Resources, Ruskin, FL) were maintained in a recirculating AHAB system (Pentair Aquatic Ecosystems, Apopka, FL) at 24-28°C on a 14:10 hour light/dark cycle at Duke University. Fish were fed twice daily with live hatched *Artemia* nauplii (90% GSL strain; Reed Mariculture, Campbell, CA) and adult zebrafish diet (388765101, Zeigler, USA). Adults (3 females, 2 males per breeder tank) were spawned for one hour and embryos collected for derivations. All procedures herein were approved by the Duke University Institutional Animal Care and Use Committee (IACUC protocol A069-22-04).

### 5.2. Germ-free Derivation and Conventionalization

Embryos were collected in sterile antibiotic solution (AB) containing 0.2-μm sterile filtered 30% Danieau’s medium (1.06 g/L NaCl, 16.3 mg/L KCl, 15 mg/L MgSO_4_, 31 mg/L Ca(NO_3_)_2_, 375 mg/L HEPES, pH 7.6), 250 ng/mL amphotericin B, 10 μg/mL kanamycin, and 100 μg/mL ampicillin. Germ-free (GF) derivation was performed at 6 hpf as previously described (13) and detailed in the Supporting Information. Groups of 25 GF embryos were reared at 28°C in 25 mL of sterile 30% Danieau’s in 50 mL 25-cm^2^ cell culture flasks (Greiner Bio-One, Monroe, NC). At 24 hpf, an 80% water change was performed with sterile Danieau’s. Sterility was confirmed by plating embryo medium on tryptic soy agar (TSA; MP Biomedicals, Irvine, CA, 1010617) plates and culturing at 28°C. Conventionalized (CV) embryos simultaneously underwent GF treatment and were subsequently conventionalized by performing an 80% water change and replacing 50% of sterile Danieau’s medium with AHAB system water at one of two timepoints: immediately following derivation at 6 hpf and at 24 hpf, to generate treatments CV6 and CV24, respectively. All CV and GF groups were maintained under consistent conditions.

### 5.3. RNA Extraction, Library Preparation, Paired-End Sequencing, and Processing

15 embryos from each CV6, CV24, or GF treatment were randomly selected at 32 hpf, pooled, and flash frozen on liquid nitrogen (experiments repeated four times). RNA was extracted from each pooled embryo sample using RNeasy® QIAGEN Mini Kit (74104, QIAGEN Inc.) according to the manufacturer’s instructions and concentration and quality was analyzed using a NanoDrop UV-Vis Spectrophotometer (Thermo Fisher Scientific, Inc., Waltham, MA). Whole-transcriptome library preparation and paired-end sequencing were performed by Duke University Sequencing and Genomic Technologies and sequences were processed as previously described (90) and included in the Supporting Information. Extracted RNA was also used for cDNA synthesis and quantitative RT PCR (qRT-PCR) to confirm relative expression of *cyp1a*, details are in the Supporting Information.

### 5.4. Pairwise Differential Gene Expression Analysis

Differential gene expression analysis was performed using R (v4.3.2) (91) and Bioconductor (v3.18) packages DESeq2 (v1.42.0) (35, 36), DEGreport (v1.38.5) (43), and org.Dr.eg.db (v3.18.0) (92) (to annotate zebrafish gene names). Pairwise comparisons were performed using the Wald test with an adjusted (adj.) p-value<0.05 and a log_2_Fold Change>|0.5| (Dataset S1). Venn diagrams were created using CRAN package ggVennDiagram (v1.2.3) (93) and volcano plots were created using Bioconductor (v3.18) package EnhancedVolcano (v1.20.0) (94). A principal component analysis (PCA) was performed among all significant pairwise DEGs and two principal components extracted. Transcript counts were normalized with a regularized log transformation and graphed with plotPCA function of DESeq2 and ggplot2 (v3.5.1) (95).

Metascape was used to screen pairwise lists of DEGs using default parameters in the custom analysis and identify significantly enriched biological process gene ontology (GO) terms (37). GO terms were considered enriched with a p-value < 0.01, a minimum count of 3, and an enrichment factor (ratio between observed : expected counts) > 1.5. A Circos plot was generated by Metascape to identify overlap between DEGs (38). Metascape was then used to group enriched GO terms into clusters (based on a Kappa score > 0.3) (96) and top terms were graphed in a heatmap. A list of all enriched GO terms is included in Dataset S2.

### 5.5. LRT Analysis of Gene Expression Trends and Enriched GO Terms

Significant DEGs were identified by the DESeq2 (v1.42.0) (35, 36) LRT function (adj. p-value<0.05, log_2_Fold Change>|0.5|), regular log transformed, and the degPatterns function of the DEGreport package (v1.38.5) (43) was used to identify gene expression trends across treatment groups using default parameters. The enrichGO function of the ClusterProfiler package (v4.11.0) (39, 40) was used to identify enriched biological process pathways using default parameters within each LRT pattern from the top 1,000 LRT DEGs (Dataset S5). LRT DEGs included in pathways related to energy metabolism, neurodevelopment, immune function, and xenobiotic metabolism were selected and RNAseq transcript counts normalized using a variance stabilizing transformation, scaled by Z-score, and graphed using CRAN package pheatmap (v1.0.12) (97).

### 5.6. Tissue-Specific Gene Set Enrichment

A tissue-specificity analysis was performed as previously described (45) to compare differential expression to 31 tissue-enriched gene sets from a published embryonic single-cell transcriptomic dataset (44). Since our dataset contained DEGs changing in different directions, we used a self-contained rotation gene set test, mroast from Limma (v3.60.3) (46) to identify significant DEGs (non-directional FDR adj. p-value<0.05) across pairwise comparisons within each tissue-enriched gene set. The number of rotations of orthogonalized residuals (‘nrot’) was set to 9999 and differential expression was corrected for false discovery rates using adjust.method = “BH”. Tissue-enriched gene sets were designated a rank based FDR adj. p-value within each pairwise comparison and graphed using CRAN packages pheatmap (v1.0.12) (97) and ggplot2 (95).

### 5.7. Proteomic Sample Preparation and Differential Protein Abundance Analysis

Fifteen embryos from each CV6, CV24, or GF treatment were selected at 32hpf, pooled, flash frozen, and stored at -80ºC until protein extraction (experiments repeated 4 times). Sample preparation and quantitative LC-MS/MS analysis was performed by Duke University Proteomics and Metabolomics Core Facility (detailed methods in the Supporting Information). Differential protein abundance analysis was performed using R (v4.3.2) (91) and Bioconductor (v3.18) package DEP (v1.26.0) (47). Data was filtered as described in the Supporting Information. Pairwise comparisons between each treatment were performed using linear models and empirical Bayes statistics. Significant differentially abundant proteins (DAPs) were identified by a p-value<0.05 and a logFold Change>|0.5| (Dataset S7). A Venn diagram of DAPs was graphed using CRAN package ggVennDiagram (v1.2.3) (93). A PCA was performed across all DAPs and graphed using plot_PCA function of DEP and ggplot (v3.5.1) (95). Significant DAPs were further analyzed using the DAVID Functional Annotation Tool (48, 49) to identify significantly enriched biological process pathways (p-value<0.05). Raw abundance counts for DAPs involved in pathways related to energy metabolism and neurodevelopmental were scaled by Z-score and graphed using CRAN package pheatmap (v1.0.12) (97). The names of DAPs graphed in heatmaps (Fig 5) are included in Dataset S8. Individual DAPs of interest were log transformed, normalized by z-score, and graphed using ggplot (v3.5.1) (95).

### 5.9. Xenobiotic Exposure

CV6, CV24, and GF embryos at 24 hpf were aqueously exposed to 10 μM benzo(a)pyrene (Sigma-Aldrich, CAS 50-32-8) in dimethyl sulfoxide (DMSO; Sigma-Aldrich, CAS 67-68-5), or a 0.1% DMSO vehicle control and raised in exposure media until endpoint collection.

### 5.10. Mitochondrial Function Assay

Mitochondrial respiration rates were measured using the Seahorse BioAnalyzer XFe96 (Agilent Technologies, Santa Clara, CA) according to previously published protocol (98, 99). Briefly, basal, maximal (using FCCP; Sigma-Aldrich, CAS 370-86-5), and non-mitochondrial (using sodium azide, Sigma-Aldrich, CAS 26628-22-8) oxygen consumption rates (OCR; pmol/O_2_/min) were measured per individual embryos. Spare mitochondrial capacity was calculated by subtracting basal OCR from maximal OCR. Embryo OCR values were normalized to the average of CV6 embryos within each experiment and graphed using GraphPad Prism 10 (v10.1.0; GraphPad Software Inc., La Jolla, CA). Two one-way ANOVAs, separated by BaP dose, with Tukey post-hoc tests, were performed using JMP® Pro (v16.2.0) (100) to determine significance (p-value<0.05). Detailed methods for mitochondrial function and copy number are in the Supporting Information.

### 5.11. Larval Behavioral Assay

Behavioral responses were measured at 5 dpf using DanioVision™ (Noldus, Wageningen, The Netherlands) as described previously (56). Briefly, total distance traveled by individual larvae was recorded across a 10-minute habituation period in the dark and 2 sequential cycles of 10-minute in the light followed by 10-minute in the dark, for a total of a 50 minutes per assay. Movement was measured in mm per minute and graphed using GraphPad Prism 10 (v10.1.0, GraphPad Software Inc., La Jolla, CA) and R package ggplot2 (v3.5.1) (95). Significance was determined using a two-way ANOVA with a Tukey post-hoc test (p-value<0.05) in JMP® Pro (v16.2.0) (100).

