## Supplemental Information for "The External Microbiome Communicates with the Developing Zebrafish (*Danio rerio*) Embryo Through the Chorion and Influences Developmental Trajectory"

### Extended Methods

**Germ-free Derivation and Conventionalization.** Germ-free derivation was performed at 6 hpf as previously described (1). Embryos were collected in sterile antibiotic solution (AB) containing 0.2-μm sterile filtered 30% Danieau's medium (1.06 g/L NaCl, 16.3 mg/L KCl, 15 mg/L $MgSO_4$, 31 mg/L $Ca(NO_3)_2$, 375 mg/L HEPES, pH 7.6), 250 ng/mL amphotericin B, 10 μg/mL kanamycin, and 100 μg/mL ampicillin. Fertilized embryos were transferred into new AB (2 embryos/1 mL) and incubated at 28°C for 4 hours. Developing embryos in the sphere stage (2) were selected for derivation, which was performed in a laminar flow hood (NuAire, Plymouth, MN). Embryos were washed three times with AB, then exposed to 0.05% polyvinylpyrrolidone for 2 minutes and 0.005% sodium hypochlorite for 20 minutes, which were each followed by three rinses with sterile 30% Danieau's. Groups of 25 germ-free (GF) embryos were transferred to 50 mL 25-$cm^2$ cell culture flasks (Greiner Bio-One, Monroe, NC) containing 25 mL of sterile 30% Danieau's and reared at 28°C. At 24 hpf, an 80% water change was performed with sterile Danieau's. Sterility was confirmed by plating embryo medium on tryptic soy agar (TSA; MP Biomedicals, Irvine, CA, 1010617) plates and culturing at 28°C.

Conventionalized (CV) embryos simultaneously underwent germ-free treatment and were subsequently conventionalized by performing an 80% water change and replacing 50% of sterile Danieau's medium with AHAB system water, creating a microbial concentration of $10^3$ colony forming units/mL (1, 3), which was confirmed by plating the conventionalized medium on TSA plates and assessing bacterial growth after 24 hours at 28°C. Conventionalization with system water was performed immediately following derivation at 6 hpf and at 24 hpf, to generate treatments CV6 and CV24, respectively. All CV and GF groups were maintained under consistent conditions throughout the duration of the experiment.

**RNA Library Preparation and Sequencing.** 200 ng RNA was used to generate KAPA mRNA HyperPrep (Roche, Basel, Switzerland) whole-transcriptome libraries. RNA was captured with magnetic oligo-dT beads and fragmented at 94°C for 2 minutes. Libraries were barcoded using Illumina Dual Index Adapters (Illumina, Inc., San Diego, CA) and amplified by PCR for 14 cycles. Amplified libraries were then pooled and the Illumina NovaSeq 6000 system (Illumina, Inc., San Diego, CA) was used for 50 base pair paired-end sequencing.

**Paired-end Sequence Processing.** Paired-end sequenced reads were processed as previously described (4) using the Duke University high-powered Computing Cluster. Illumina TruSeq3-PE adapters and low quality reads (Q<20) were trimmed using Trimomatic v0.39 (5). The zebrafish reference genome was downloaded from Ensembl (GRCz11, Ensembl 110, GCA_000002035.4) (6) and extracted splice sites and exons were used to create an index using HISAT2 v2.2.0 (7). Trimmed paired-end reads were mapped and aligned to the indexed zebrafish reference genome using HISAT2. Generated SAM files were then sorted, converted to BAM format, and sequencing lanes merged using Samtools v1.10 (8). Merged alignments were assembled into full-length transcripts and quantified using StringTie v2.2.1 (9). The Python scriplet 'prepDE.py3' (http://ccb.jhu.edu/software/stringtie/dl/prepDE.py3) with the '-e' option was used to generate readable counts of transcript abundances. Transcript counts were filtered to include only those with a total >10 reads.

**cDNA Synthesis and quantitative RT PCR (qRT-PCR).** CV and GF embryos were generated using the methods defined in Section 5.2 of the main text, including additional time points of conventionalization: 6 hpf (CV6), 12 hpf (CV12), 24 hpf (CV24), and 30 hpf (CV30). 15 embryos from each treatment were randomly selected at 32 hpf, pooled, flash frozen, and stored at -80°C. All experiments were repeated 3 times. RNA was extracted from each pooled sample using RNeasy® QIAGEN Mini Kit (74104, QIAGEN Inc.) according to the manufacturer's instructions. RNA concentration and quality was confirmed using a NanoDrop UV-Vis Spectrophotometer (Thermo Fisher Scientific, Inc., Waltham, MA).

Extracted RNA was diluted to 150 ng/μl and 15 μl per sample were added to a well containing 2 μl 10x RT buffer, 0.8 μl 100 mM dNTPs, 2 μl 10x random primers, 1 μl reverse transcriptase (MultiScribe), 1 μl RNase inhibitor, and 3.2 μl nuclease-free water (4374966, Applied Biosystems™, Waltham, MA). cDNA was generated on a PTC-200 Gradient Thermal Cycler (MJ

Research Inc., Quebec, Canada) under the following reaction conditions: 25°C for 10 minutes, 37°C for 2 hours, and 85°C for 5 minutes, and stored at -80°C. cDNA was diluted to 3 ng/µl and 2 µl per sample were added to a well containing 10 µl Luna Universal Probe qPCR Master Mix (M3004, New England Biolabs, Ipswich, MA), 1.6 µl forward/reverse *cyp1a* primer, 0.4 µl *cyp1a* probe, 0.8 µl forward/reverse *actb1* primer, 0.2 µl *actb1* probe, and 5 µl nuclease-free water. Primers and probes were provided by IDT (Integrated DNA Technologies, Inc., Morrisville, NC), sequence properties and off-target compatibility was assessed using SciTools™ OligoAnalyzer™ Tool (10), and sequences are included in Table S1.

Transcript abundance was quantified via reverse transcription qPCR (RT-qPCR) on an Analytik Jena qTower$^3$ (Analytik Jena, Germany) under the following reaction conditions: 95°C for 120 seconds, with 45 cycles of 95°C for 15 seconds and 52°C for 60 seconds. *Actb1* was used as the housekeeping reference gene and relative transcript abundance of *cyp1a* was quantified using the $\Delta\Delta$Ct method (11) and the $\Delta$Ct method, normalizing to the average of CV6 replicates, and relative expression for each method graphed using ggplot (v3.5.1) (12). Primer efficiencies were calculated at 103% and 109% for *cyp1a* and *actb1*, respectfully.

**Proteomic Sample Preparation and Quantitative LC-MS/MS Analysis.** Samples were probe sonicated in 5% SDS and total protein content was quantified using a Pierce™ Detergent Compatible Bradford Assay Kit (ThermoFisherScientific, Waltham, MA). 20 µg of extracted protein from each sample was reduced with 10 mM DTT at 80°C for 10 minutes, alkylated with 25 mM iodoacetamide for 30 minutes at 21ºC, and 1.2% phosphoric acid and 200 µl of S-Trap (Protifi) binding buffer (90% MeOH/100mM TEAB) were added to each sample for denaturation. Proteins were bound to the S-Trap micro cartridge, digested with 80 ng/µL trypsin (Promega) at 47°C for 1 hour, and eluted by adding 50 mM TEAB, 0.2% FA, and 50% ACN/0.2% FA. Samples were lyophilized and resuspended in 20 µl of 1% TFA/2% MeCN. An equal volume of each sample was combined to produce a quality control for the study design.

1 µl of each sample was analyzed in a randomized order by Quantitative LC-MS/MS on a Vanquish Neo UHPLC coupled to a Thermo Orbitrap Astral (ThermoFisherScientific, Waltham, MA). Samples were bound to a PepMap C18 0.3 mm x 5 mm column and separated with a 150 µm ID x 8 cm, 1.5 µm (PepSep) column using increasing concentration of 5 to 30% acetonitrile with 0.1% formic acid for 30 minutes at 55 °C. Data was retrieved using a data-independent acquisition (DIA) mode. DIA data were converted to .htrms format and processed in Spectronaut 18 (18.7.240325.55695 Biognosys AG, Schlieren, Switzerland). A spectral library was built for zebrafish using a NCBI database and a tryptic specificity up to 2. Protein groups were filtered by a 1% peptide/protein false-discovery rate, used for local normalization (13), and protein abundances were calculated using the MaxLFQ algorithm (14).

**Proteomic Data Filtration and Normalization.** Bioconductor (v3.18) package DEP (v1.26.0) (15) was used to filter proteomic data for proteins identified in at least three replicates of at least one treatment using a threshold value of 1 in the filter_missval function. Normalized data was compared to non-normalized data, which revealed a variance stabilizing transformation produced right-skewed dataset (Fig. S12). Therefore, the non-normalized filtered data that followed a Gaussian distribution was used for further analysis. A comparison of all significant proteins (adj. p-value<0.05) for normalized and non-normalized data are included in Fig. S13. Missing values were random (MAR) and a k-nearest neighbor approach (rowmax=0.9) was used to impute missing values.

**Mitochondrial Function Assay.** Mitochondrial respiration rates were measured using the Seahorse BioAnalyzer XFe96 (Agilent Technologies, Santa Clara, CA) according to previously published protocol (16, 17). Non-deformed embryos were randomly selected for assay and placed in the bottom, center of individual spheroid chambers containing 150 µl of 65 ppm ASW. Oxygen consumption rate (OCR; pmol/$O_2$/min) was measured per individual embryos at basal levels and following the injection of electron transport chain inhibitors. Maximal mitochondrial respiration was measured following the injection of 6 µM carbonyl cyanide-p-trifluoromethoxyphenylhydrazone (FCCP; Sigma-Aldrich, CAS 370-86-5) and non-mitochondrial respiration was measured following injection of 6.25 mM sodium azide (Sigma- Aldrich, CAS 26628-22-8). Basal OCR was measured for 8 cycles, post-injection of FCCP was measured for 8 cycles, and post-injection of sodium azide was measured for 35 cycles, with a 1-minute mix, 1-minute wait, and 2-minute measure per cycle. Assays were repeated across three experiments.

Following each run, non-mitochondrial oxygen consumption rate (OCR) was identified as the average of the lowest three OCR measurements after sodium azide injection, basal OCR was calculated as the average of the lowest three OCR measurements prior to FCCP injection minus non-mitochondrial OCR, maximal OCR was the average of the three highest measurements following FCCP injection minus non-mitochondrial OCR, and spare capacity was calculated by subtracting basal OCR from maximal OCR. Embryo OCR values were normalized to the average of CV6 embryos within each experiment and graphed using GraphPad Prism 10 (v10.1.0; GraphPad Software Inc., La Jolla, CA). Values that fell outside ±1.5(IQR) based on a standard normal distribution were excluded from statistical analysis if removal did not significantly skew the sample distribution and change results. Two one-way ANOVAs, separated by BaP dose, with Tukey post-hoc tests, were performed using JMP® Pro (v16.2.0) (18) to determine significance (p-value<0.05) in average OCR values.

**Mitochondrial Copy Number Assay.** Mitochondrial copy number was measured as previously described (19). Briefly, 15 embryos from GF and CV24 treatments were pooled at 48hpf, with 3 experimental replicates, flash frozen, and DNA extracted using DNeasy blood and tissue kit (QIAGEN, Germany). 2 µl of DNA (3 ng/µl) from each sample was added in triplicate to individual wells of a 96-well plate with 12.5 µl Power SYBR™ Green PCR Master Mix (4368577, Thermo Fisher Scientific, Waltham, MA), 1 µl of forward primer, 1 µl of reverse primer, and 8.5 µl of nuclease-free water. Mitochondrial DNA was detected by the *mt-nd1* region of the mitochondrial genome and normalized to nuclear DNA (*actb1*). For each sample, detection of mitochondrial and nuclear DNA was performed in separate wells. Primer sequences can be found in Table S2. Primers were provided by IDT (Integrated DNA Technologies, Inc., Morrisville, NC) and sequence properties and off-target compatibility was assessed using SciTools™ OligoAnalyzer™ Tool (10). Gene abundance was quantified via quantitative-PCR (qPCR) on an Analytik Jena qTower$^3$ (Analytik Jena, Germany) under the following reaction conditions: 50°C for 120 seconds, 95°C for 10 minutes, 40 cycles of 95°C for 15 seconds and 60°C for 60 seconds, followed by a melt curve. Relative abundance of *mt nd-1* was quantified using the ΔΔCt method (11) and relative expression for each method graphed using GraphPad Prism 10 (v10.1.0; , GraphPad Software Inc., La Jolla, CA). A one-way ANOVA was performed using JMP® Pro (v16.2.0) (18) to determine significance (p-value<0.05).

**Larval Behavioral Assay.** Behavioral responses were measured at 5 dpf using DanioVision™ (Noldus, Wageningen, The Netherlands). Non-deformed larvae were randomly selected and transferred into individual wells of a clear, flat-bottom 96-well plate (Greiner Bio-One, Monroe, NC) with approximately 200 µL of sterile (GF) or conventionalized (CV6 and CV24) 30% Danieau's. Assays were repeated across three experiments. Larval location in each well was recorded every 0.016 seconds using EthoVision XT14 software (Noldus, Wageningen The Netherlands) during a 10-minute habituation period in the dark, followed by 2 cycles of 10-minute light followed by 10-minute dark, for a total of a 50 minutes per assay. Total distance traveled was measured by mm/minute based on the recorded larval location during each phase and graphed using GraphPad Prism 10 (v10.1.0, GraphPad Software Inc., La Jolla, CA) or R package ggplot2 (v3.5.1) (12). Values that fell outside ±1.5(IQR) based on a standard normal distribution were excluded from data analysis and the remaining individuals were averaged within treatments. Significance was determined using a two-way ANOVA with a Tukey post-hoc test (p-value<0.05) in JMP® Pro (v16.2.0) (18).

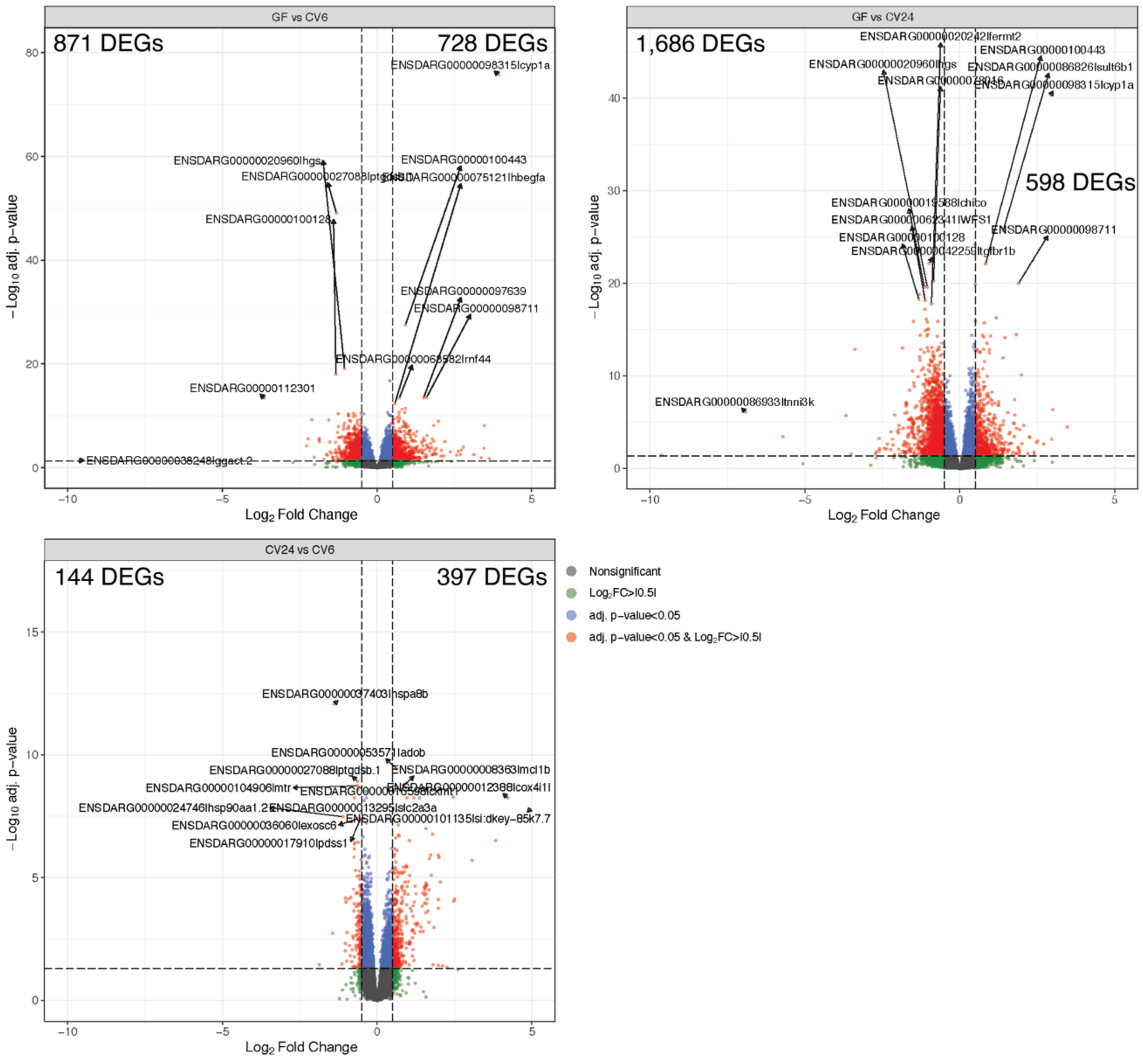


**Fig. S1.** Significant DEGs from DESeq2 pairwise comparisons (adj. p-value<0.05 and $\log_2$Fold Change>|0.5|). Volcano plots were created using Bioconductor (v3.18) package EnhancedVolcano (v1.20.0) (20). Each point represents the average value for a gene transcript across all 4 replicates (each replicate with 15 pooled embryos) and are plotted by $\log_2$Fold Change (x-axis) and -$\log_{10}$ adj. p-value (y-axis). Thresholds are drawn at a $\log_2$Fold Change>|0.5| and adj. p-value<0.05. Transcripts that fall within these significance bounds are denoted as follows: grey (nonsignificant DEGs), green ($\log_2$Fold Change>|0.5|), blue (adj. p-value<0.05), and red (adj. p-value<0.05 and $\log_2$Fold Change>|0.5|). The number of DEGs with a negative and positive $\log_2$Fold Change are listed for each pairwise comparison.

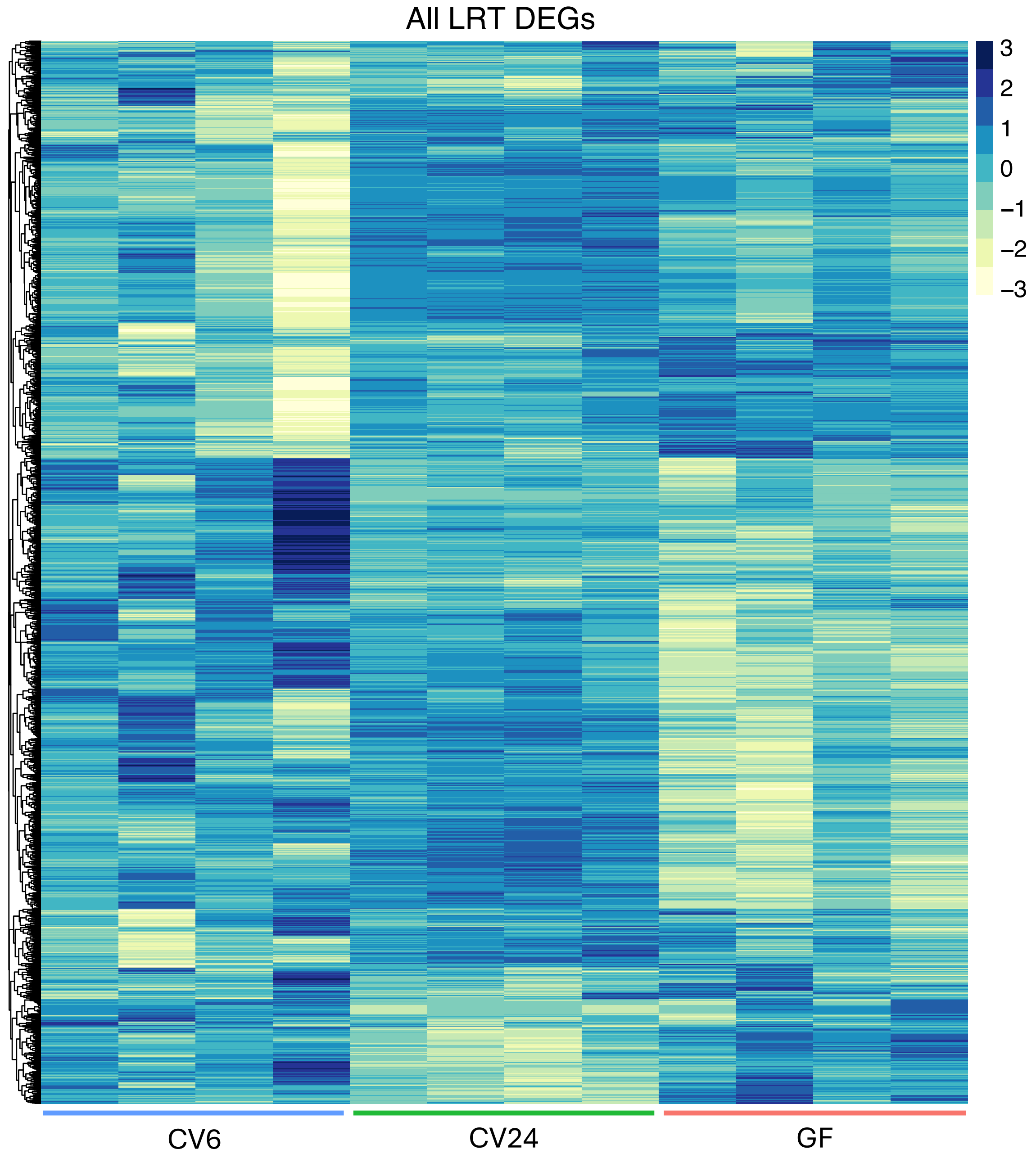


**Fig. S2.** Heatmap of normalized transcript counts for all 1,769 significant LRT DEGs (adj. p-value<0.05, $log_2$Fold Change>|0.5|) graphed using CRAN package pheatmap (v1.0.12) (21). Transcript counts for each of the 4 sequenced sample replicates per treatment are shown on the x-axis, with each replicate represented by 15 pooled embryos. Counts are scaled by Z-score within rows and gene identifiers are shown on the y-axis.

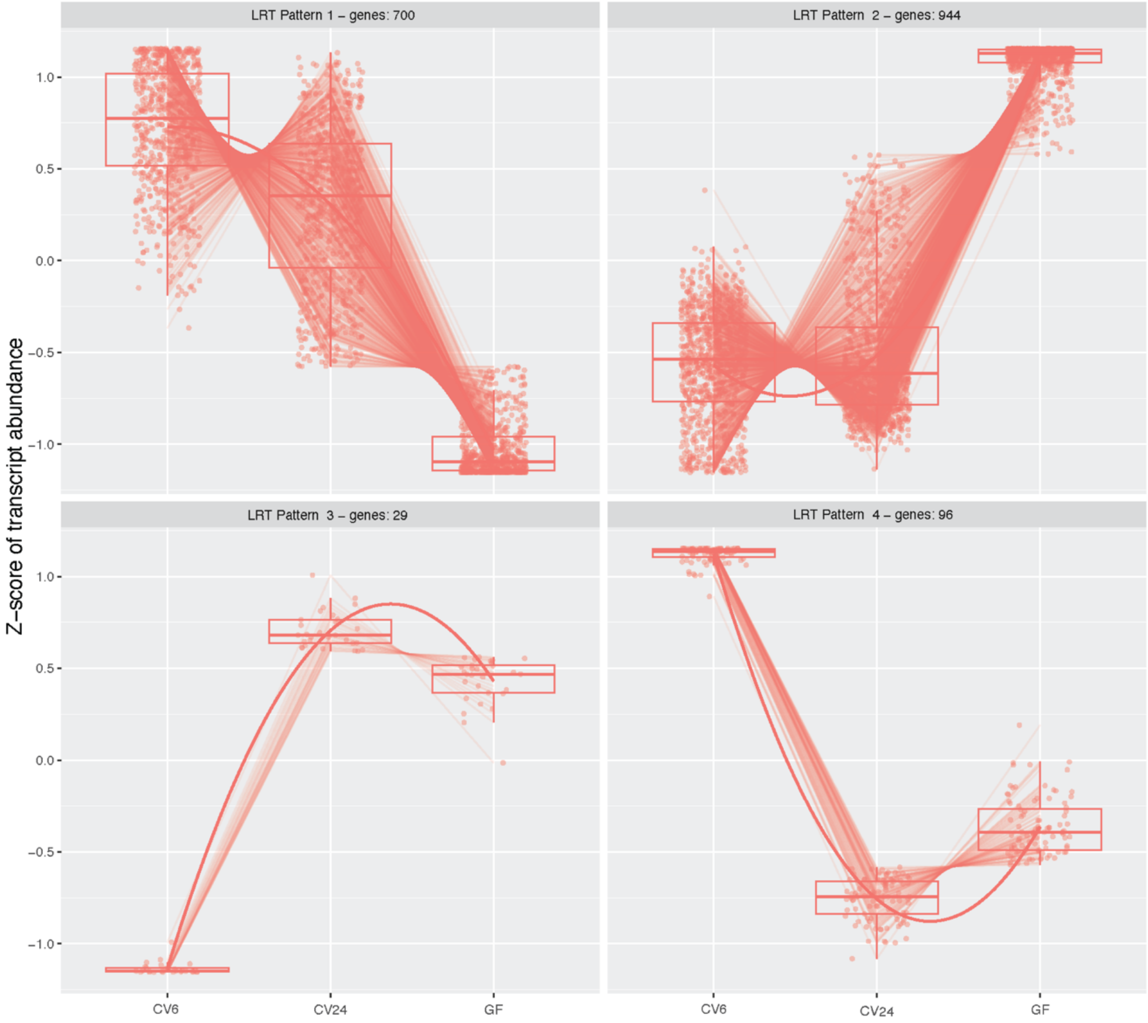


**Fig. S3.** LRT gene expression patterns identified by the degPatterns function of the DEGreport package (v1.38.5) (22) within all 1769 significant DEGs identified by the DESeq2 LRT command (23, 24) across treatments (adj. p-value<0.05, $log_2$Fold Change>|0.5|). Each group illustrates the number of genes with a similar expression pattern identified using DESeq2 DEGpatterns. Each dot represents the mean transcript abundance across all 4 replicates within a treatment for a particular gene and mean transcript counts are scaled by Z-score. Data are displayed by treatment (x-axis) and Z-score of transcript abundance (y-axis).

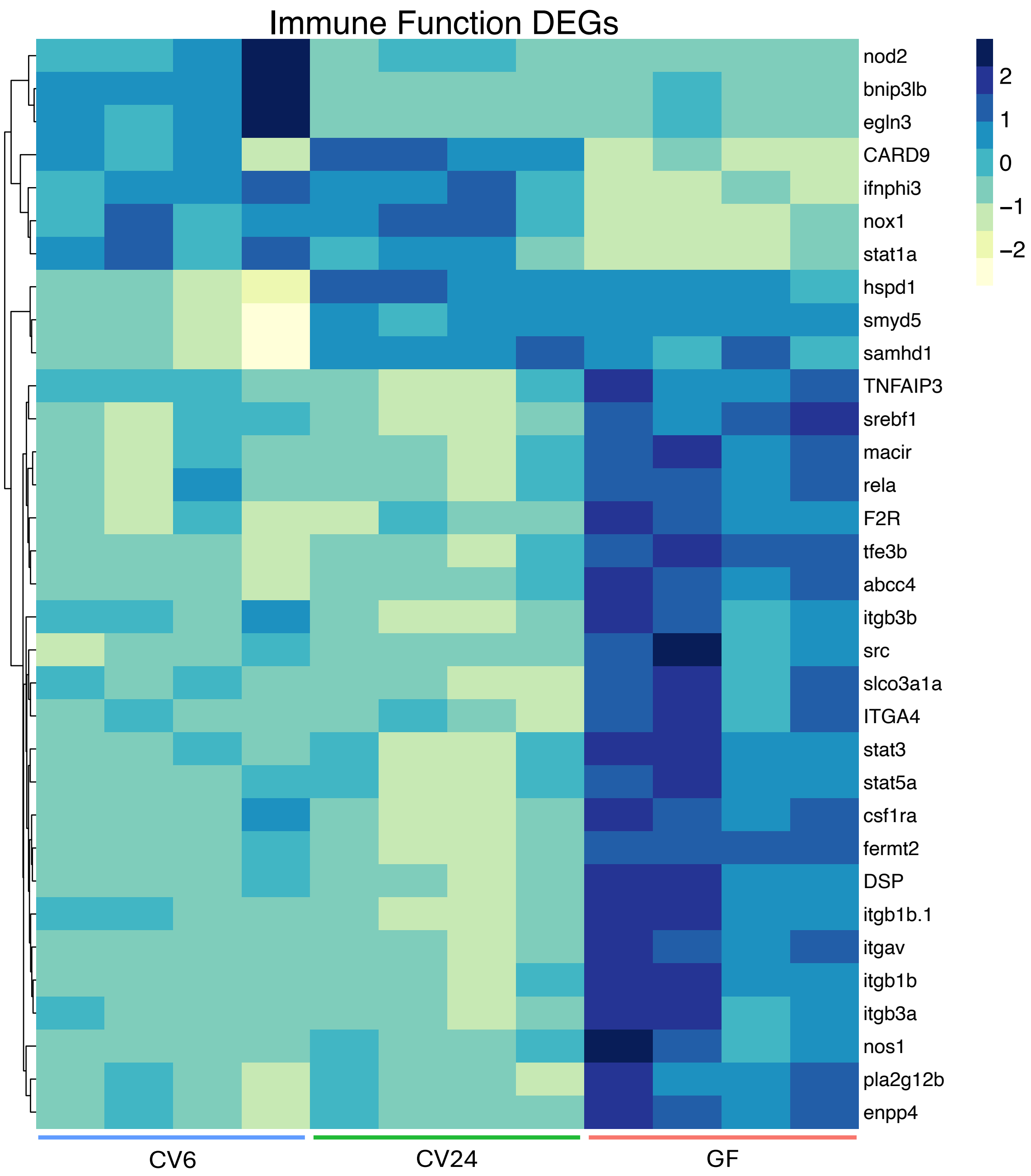


**Fig. S4.** Heatmap of normalized transcript counts for DEGs involved in immune function graphed using CRAN package pheatmap (v1.0.12) (21). DEGs represent all genes included in significantly enriched biological process GO terms (p-value<0.05) and among the top 1000 DEGs identified by the LRT function from the DESeq2 package (adj. p-value<0.05, $\log_2$Fold Change>|0.5|) (23–26). Transcript counts for each of the 4 sequenced sample replicates per treatment are shown along the x-axis. Each sample is represented by 15 pooled embryos. Counts are scaled by Z-score within rows and gene identifiers are shown on the y-axis.

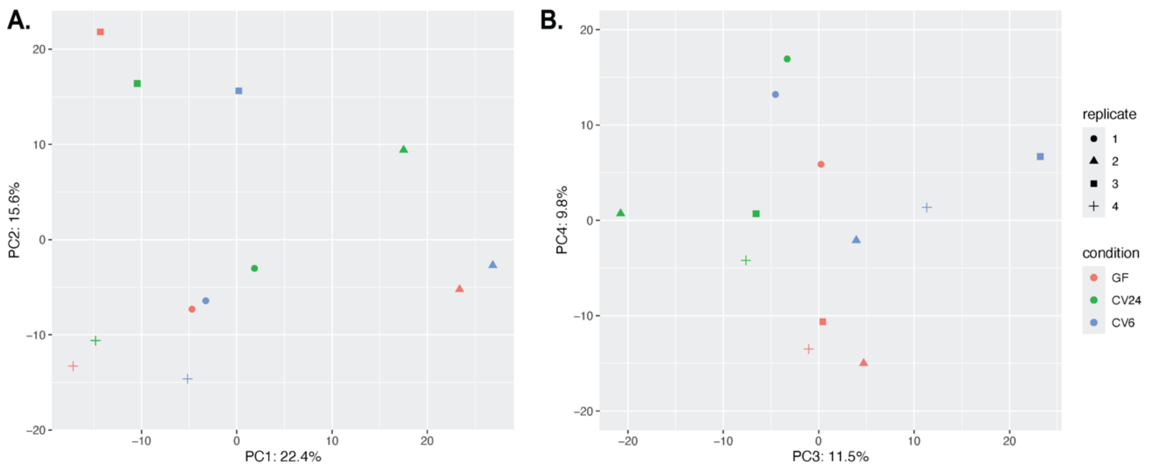


**Fig. S5.** Principal component analysis (PCA) plots of non-normalized differentially expressed protein counts graphed with plot_PCA function of DEP (15) using default parameters and ggplot (v3.5.1) (12). **(A)** PCs 1 and 2 for all proteins included in analysis after filtering thresholds set to 'thr = 1' using the 'filter_missval' command of Bioconductor DEP package (15). **(B)** PCs 3 and 4 for all proteins included in analysis after filtering thresholds set to 'thr = 1' using the 'filter_missval' command of Bioconductor DEP package (15). Each point represents a quantified sample containing 15 pooled embryos at 32hpf. Each condition is represented by 4 sequenced replicates and labeled by experimental replicate (shape) and treatment condition: CV6 (blue), CV24 (green), and GF (pink). PC1 and PC2 suggest experimental replicate is one of the major factors explaining the variance described by the first two principal components, while treatment condition is one of the major factors explaining the variance described by PC3 and PC4. However, given the batch effect demonstrated by the strong clustering by replicate in PC1 and PC2, the observed differences in protein quantification between treatments must be robust enough to overcome the observed similarities within replicates. While replicates 1 and 2 were performed in parallel on the same day, similarly to replicates 3 and 4, each replicate used embryos derived from different sets of parents. Since distinct clustering between replicates 1 and 2 or between 3 and 4 were not observed, we suggest the observed clustering within replicates is not attributed to an experimental batch effect, but rather the distinct genetic makeup of each set of embryos, although further testing is needed to confirm protein expression remains consistent upon subsequent breedings from the same parents.

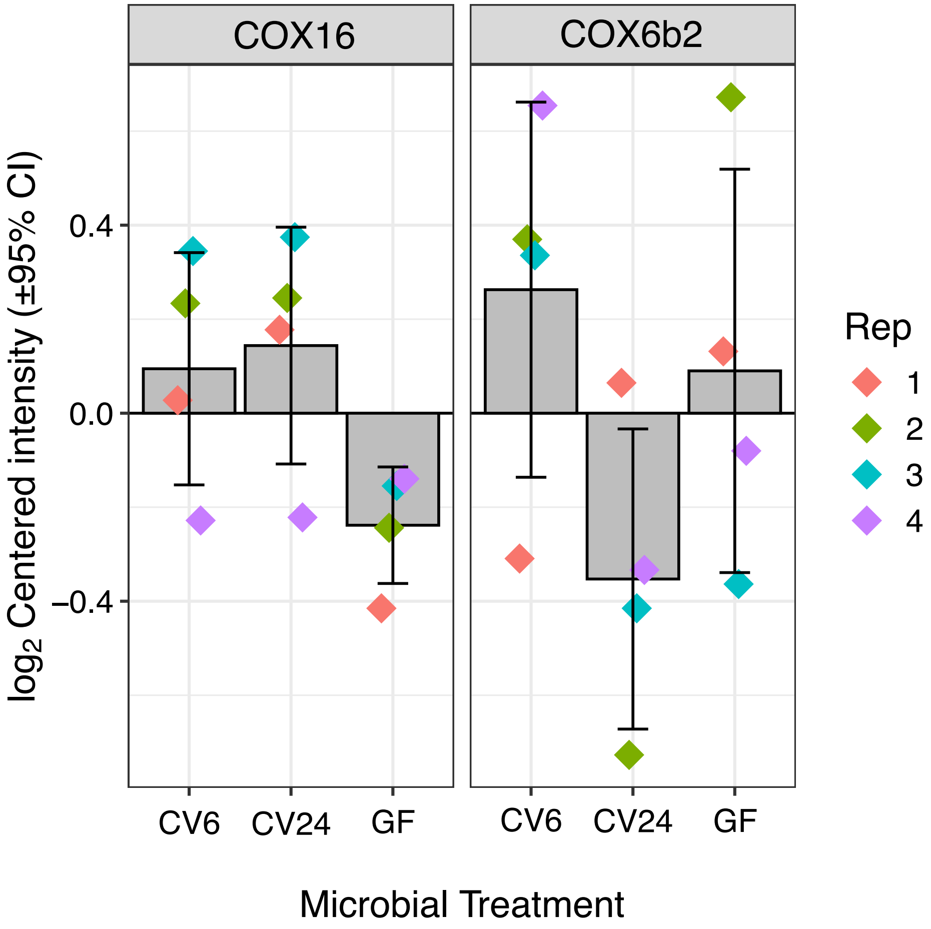


**Fig. S6.** Cytochrome c oxidase assembly factor (COX16) and cytochrome c oxidase subunit 6B2 (COX6b2) protein expression in CV6, CV24, and GF embryos at 32hpf. Each point represents a quantified sample containing 15 pooled embryos. Replicates are labeled by color: 1 (pink), 2 (green), 3 (blue), and 4 (purple). Data is displayed by microbial treatment (x-axis) and $log_2$Centered intensity (±95% Confidence Interval) (y-axis).

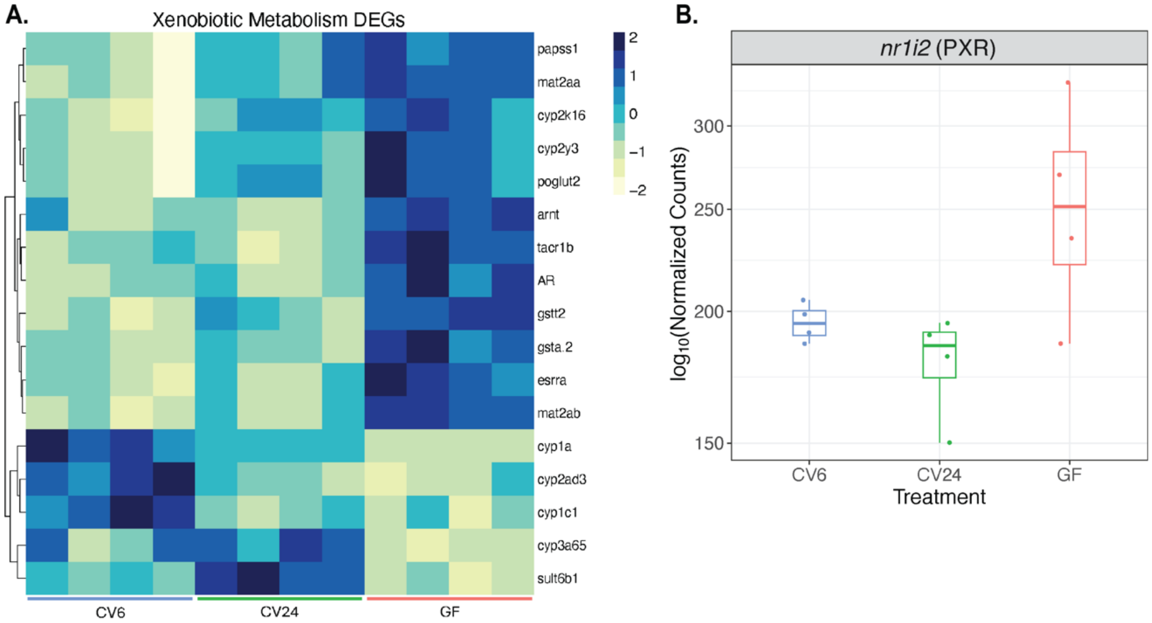


**Fig. S7. (A)** Heatmap of normalized transcript counts for DEGs involved in xenobiotic metabolism graphed using CRAN package pheatmap (v1.0.12) (21). DEGs represent all genes included in significantly enriched biological process GO terms (p-value<0.05) and among the top 1000 DEGs identified by the LRT function from the DESeq2 package (adj. p-value<0.05, $\log_2$Fold Change>|0.5|) (23–26). Transcript counts for each of the 4 sequenced sample replicates per treatment are shown along the x-axis. Each sample is represented by 15 pooled embryos. Counts are scaled by Z-score within rows and gene identifiers are shown on the y-axis. **(B)** $\log_{10}$ Normalized transcript count for PXR (*nr1i2*) across CV6, CV24, and GF groups graphed using ggplot (v3.5.1) (12). Each point represents a sequenced sample containing 15 pooled embryos at 32 hpf. Each condition is represented by 4 sequenced replicates and labeled by treatment condition for CV6 (blue), CV24 (green), and GF (pink). Data are displayed by microbial treatment (x-axis) and $\log_{10}$(Normalized Counts) (y-axis).

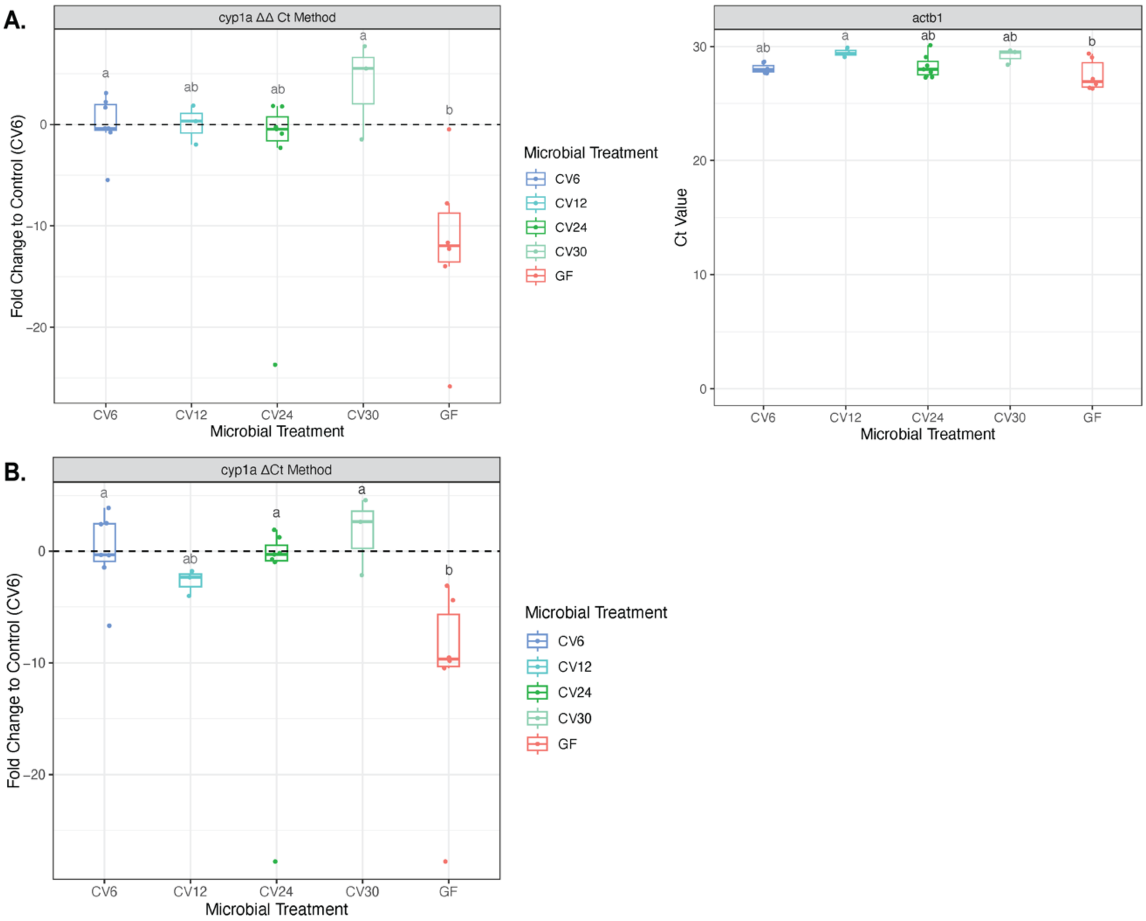


**Fig. S8.** Fold change in *cyp1a* gene expression in CV6, CV12, CV24, CV30, and GF embryos at 32hpf relative to the average expression of CV6 embryos graphed using ggplot (v3.5.1) (12). **(A)** Relative expression calculated using the ΔΔCt method (11) based on the relative expression of actin-β1 (*actb1*) and CV6 as the control average and **(B)** Relative expression calculated using the ΔCt method based on CV6 as the control average. Each point represents a sequenced sample containing 15 pooled embryos at 32hpf. CV12 and CV30 groups are represented by 3 pooled replicates and CV6, CV24, and GF groups are represented by an additional 3-4 replicates to confirm expression of the samples used for RNA sequencing. For groups that had more than 5 replicates, values that fell outside ±1.5(IQR) based on a standard normal distribution were excluded from data analysis and the remaining individuals within treatments were averaged. Significance determined by 1-way ANOVA, Tukey's HSD, $p<0.05$. Treatments are labeled by condition for CV6 (dark blue), CV12 (light blue), CV24 (dark green), CV30 (light green), and GF (pink). Data is displayed by microbial treatment (x-axis) and Fold change to Control (CV6) (y-axis). A dotted line is placed at 0 to represent the fold change of each sample in relation to the average expression of CV6 samples. The Ct values for *actb1* in each sample are shown to illustrate slight variation in gene expression in the house keeping gene across samples. Since minor differences were noted in *actb1* expression across treatment groups, an equal amount of RNA (6ng) was added to each reaction and the relative expression was calculated without a house keeping gene, using the ΔCt method based on the average Ct value of CV6 as the control. Both methods confirm the same trend in relative expression of *cyp1a* across treatments.

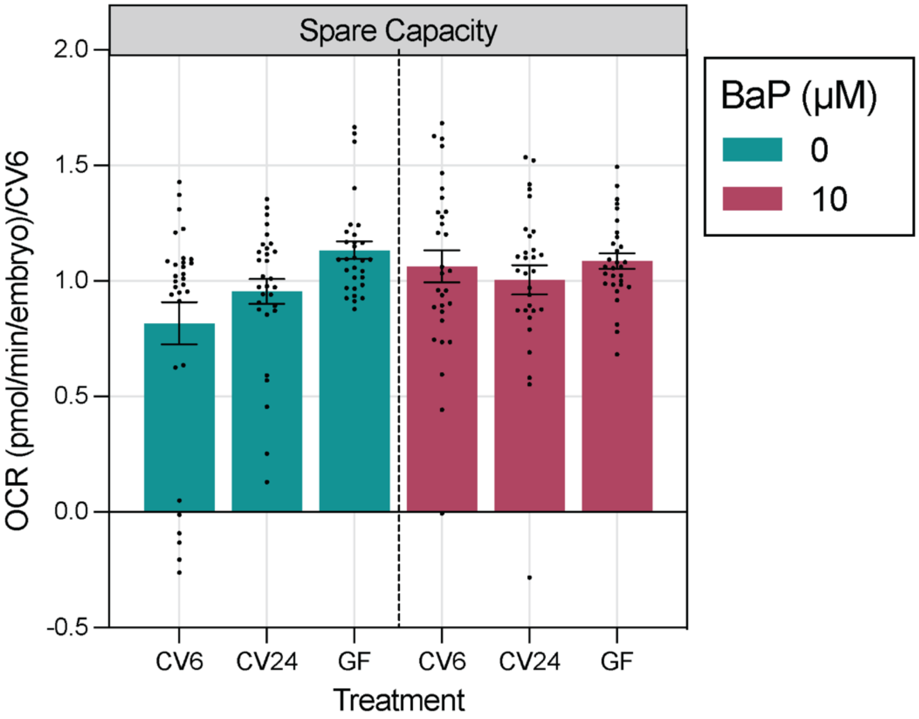


**Fig. S9.** Spare respiratory capacity in CV6, CV24, and GF embryos at 32hpf dosed with either a vehicle control (0.1% DMSO) or 10µM BaP. Data normalized to the average of CV6 embryos across replicates. No statistical significance found by microbial treatment or BaP exposure. Significance determined by two 1-way ANOVAs, separated by BaP treatment, Tukey's HSD, $p<0.05$ using JMP® Pro (v16.2.0) (18). Error bars represent standard error mean. n=30-44. Data is displayed by microbial treatment (x-axis), grouped by BaP treatment [0µM BaP (blue) and 10µM BaP (red)], and OCR (pmol/min/embryo), normalized to CV6 average (y-axis) and graphed using GraphPad Prism 10 (v10.1.0; , GraphPad Software Inc., La Jolla, CA).

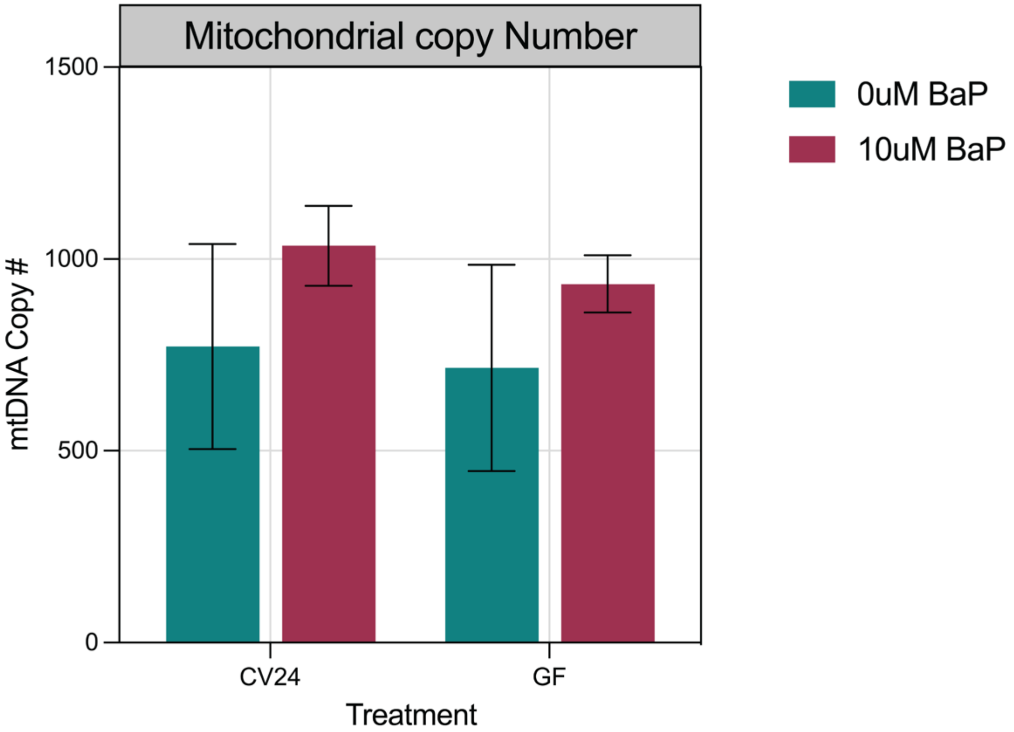


**Fig. S10.** Mitochondrial (mtDNA) copy number in CV24 and GF larvae at 48hpf, prior to embryonic hatching, dosed with either a vehicle control (0μM BaP; 0.1% DMSO) or 10μM BaP. Each bar represents the average relative mtDNA copy number identified from 3 replicates of 15 pooled embryos. mtDNA copy number determined from sample DNA by qPCR compared to the quantity of *actb1*, as the HK gene using methods previously described (11, 19). No statistical significance found by microbial treatment or BaP exposure. Significance determined by a 2-way ANOVA, Tukey's HSD, $p<0.05$ using JMP® Pro (v16.2.0) (18). Error bars represent standard error mean. Data is displayed by microbial treatment (x-axis), grouped by BaP treatment [0μM BaP (blue) and 10μM BaP (red)], and mtDNA copy # (y-axis) and graphed using GraphPad Prism 10 (v10.1.0; , GraphPad Software Inc., La Jolla, CA).

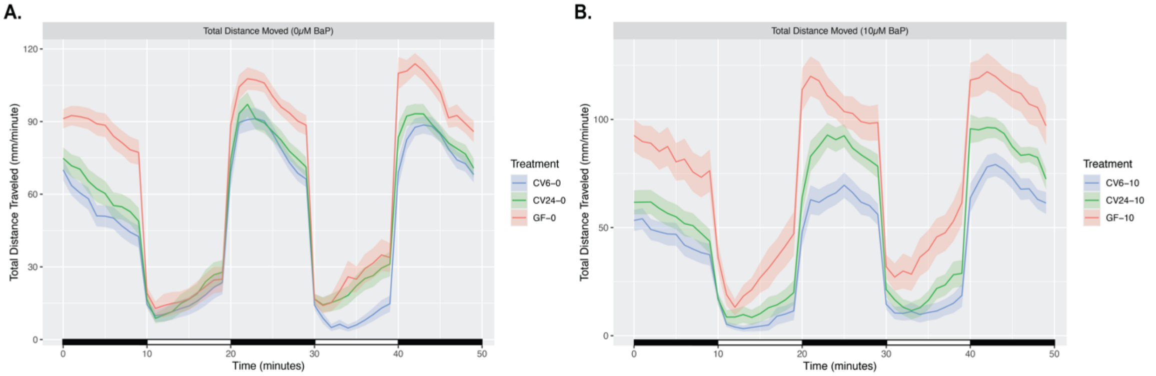


**Fig. S11.** Total distance moved across time (0-9 min – habituation, 10-19 min – light, 20-29 min – dark, 30-39 min – light, 40-49 min – dark) in CV6, CV24, and GF larvae at 5dpf dosed with either a vehicle control (0μM BaP; 0.1% DMSO) or 10μM BaP. n=35-64. Larval location was tracked and recorded every 0.016 seconds using DanioVision$^{TM}$ and EthoVision XT14 software (Noldus, Wageningen The Netherlands). Data is displayed by time (minutes) (x-axis) and total distance traveled (mm/minute) (y-axis) and graphed using ggplot (v3.5.1) (12). Each condition is labeled by CV6 (blue), CV24 (green), GF (pink), and separated by **(A)** 0μM and **(B)** 10μM BaP. Error intervals represent standard error.

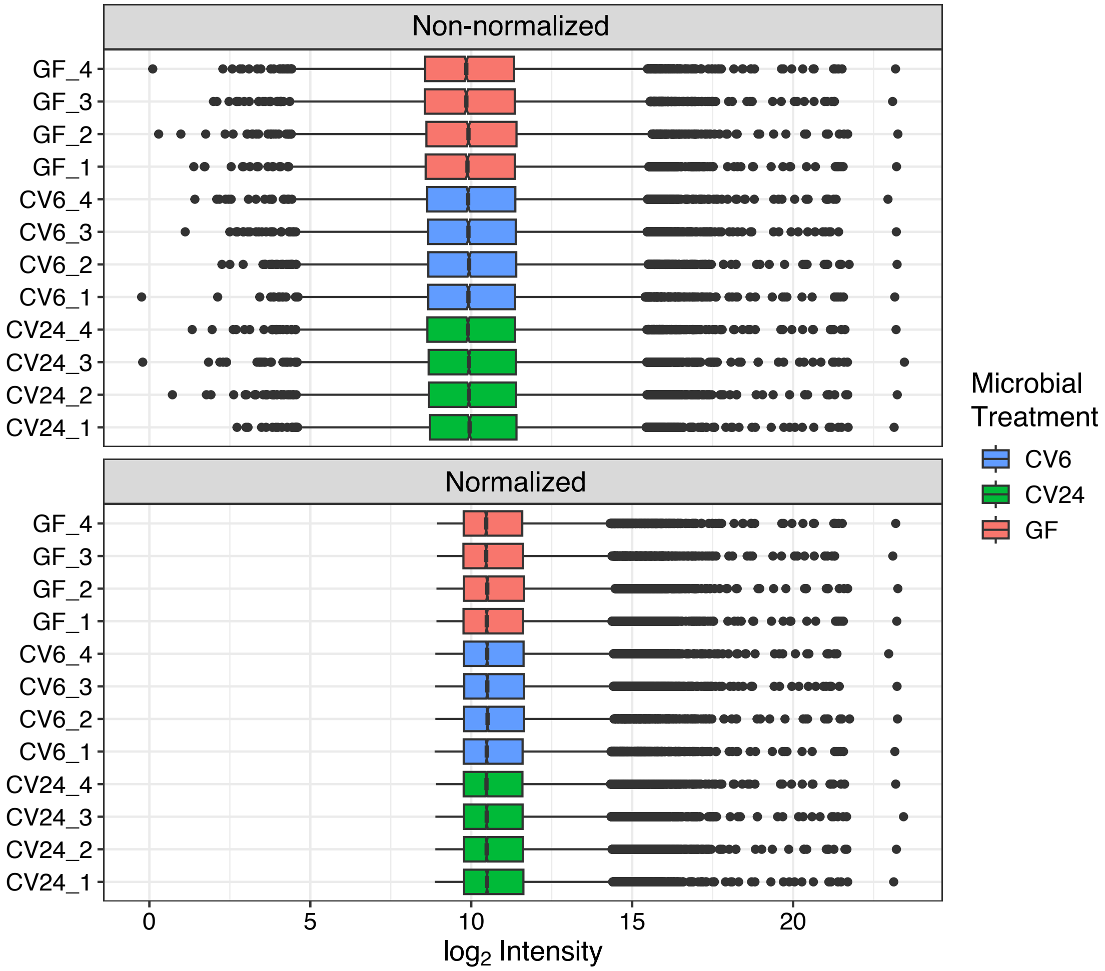


**Fig. S12.** Comparison of non-normalized and normalized filtered protein expression data, analyzed using Bioconductor (v3.18) package DEP (v1.26.0) (15). Box plots show the effect of normalization on the data for each sample, represented by 15 pooled embryos. Normalization removes low intensity counts. Data is displayed by $log_2$Intensity (x-axis) and sample ID (y-axis). Treatments are labeled by condition: CV6 (blue), CV24 (green), and GF (pink).

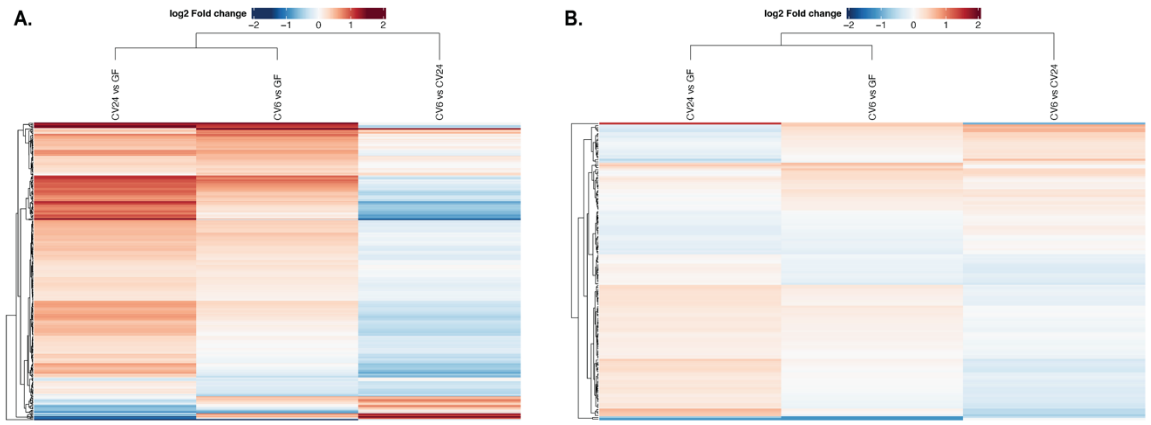


**Fig. S13.** Comparison of the fold change across tested pairwise comparisons for **(A)** non-normalized and **(B)** normalized data. Fold change is quantified as the difference between the average of treatments analyzed in each pairwise comparison using Bioconductor (v3.18) package DEP (v1.26.0) (15). Significantly differentially expressed proteins determined by adjusted [adj.] p-value<0.05. Protein counts are scaled by Z-score within rows and clustered by hierarchal clustering.

**Table S1.** Primer and probe sequences for *cyp1a* and *actb1* (housekeeping reference gene), associated melting temperature and percent GC content. Probe for *actb1* includes locked nucleic acid nucleotides to increase melting temperature, specificity, and stability. Locked nucleic acid nucleotides are preceded by a '+'.

| Ensembl Gene ID | Gene Name | Primer Type | Sequence (5'-3') | Tm(°C) | GC(%) |
|---|---|---|---|---|---|
| ENSDARG00000098315 | cyp1a | Forward | CCACTGCGAAGACCGAAAAC | 56.6 | 55.0 |
| | | Probe (SUN ZEN / Iowa Black™ FQ) | 5SUN/AACTCCAAC/ZEN/CTGCAAGTGTCCGAT/3IABkFQ | 60.7 | 50.0 |
| | | Reverse | AGATAGACAACCGCCCAGGA | 57.9 | 55.0 |
| ENSDARG00000037746 | actb1 | Forward | CCTGAATCCCAAAGCCAACA | 55.6 | 50.0 |
| | | Probe (FAM ZEN / Iowa Black™ FQ) | 56FAM/AGAG+AA+GAT+GACA+CA+GATC/3IABkFQ | 59.2 | 42.1 |
| | | Reverse | GGAAGAGCGTAACCCTCATAGA | 55.8 | 50.0 |

**Table S2.** Primer sequences for *mt nd-1* and *actb1* (housekeeping reference gene), associated melting temperature and percent GC content. Primers used to quantify mitochondrial copy number.

| Ensembl Gene ID | Gene Name | Primer Type | Sequence (5'-3') | Tm(°C) | GC(%) |
|---|---|---|---|---|---|
| ENSDARG00000063895 | mt-nd1 | Forward | CGTTTACCCCAGATGCACCT | 64.2 | 55.0 |
| | | Reverse | GTGCGATTGGTAGGGCGATA | 63.9 | 55.0 |
| ENSDARG00000037746 | actb1 | Forward | TGGATACCTGACCGAGAGCT | 64.1 | 55.0 |
| | | Reverse | AGACAACTCTTACGGCTGGC | 64.0 | 55.0 |

**Dataset S1 (separate file).** Significant DEGs (adj. p-value<0.05, $log_2$Fold Change>|0.5|) identified across pairwise comparisons by the Wald Test using DESeq2 (23, 24).

**Dataset S2 (separate file).** Enriched biological process gene ontology (GO) terms and associated genes identified within each pairwise comparisons using the multi-list input in Metascape (27). Pathways with a p-value < 0.01, a minimum count of 3, and an enrichment factor (ratio between observed:expected counts) > 1.5 were considered significant and included in analysis.

**Dataset S3 (separate file).** Significant DEGs (adj. p-value<0.05, $log_2$Fold Change>|0.5|) identified by a likelihood ratio test using the DESeq2 LRT function (23, 24).

**Dataset S4 (separate file).** DEGs included in each gene expression pattern within the top 1000 DEGs identified by the DESeq2 LRT command across all treatments (adj. p-value<0.05, $log_2$Fold Change>|0.64|). Each tab lists all DEGs genes from a different pattern, as displayed in Figure 2.

**Dataset S5 (separate file).** Enriched biological process gene ontology (GO) terms and associated genes identified within each gene expression cluster using the enrichGO function of the ClusterProfiler package (v4.11.0) (25, 26). Pathways with a p-value<0.05 were considered significant and included in analysis. GO terms and included DEGs for each gene expression cluster are included on separate tabs.

**Dataset S6 (separate file).** Normalized DEGs involved in energy metabolism and neurodevelopmental pathways included in heatmaps shown in Figure 3. DEGs identified among gene expression clusters using the enrichGO function of the ClusterProfiler package (v4.11.0) (25, 26). Pathways with a p-value<0.05 were considered significant and included in analysis. DEGs for energy metabolism and neurodevelopment are included on separate tabs and columns show individual normalized transcript counts for each sample replicate.

**Dataset S7 (separate file).** Significant differentially abundant proteins (DAPs; p-value<0.05, $log_2$Fold Change>|0.5|) identified across pairwise comparisons using Bioconductor (v3.18) package DEP (v1.26.0) (15). To identify DAPs, non-normalized data was used, a threshold value of 1 was set in the filter_missval command, and a k-nearest neighbor approach with a rowmax of 0.9 was used to impute missing values. Proteins significant for each pairwise comparisons are included in a separate tab.

**Dataset S8 (separate file).** Significant DAPs (p-value<0.05) involved in energy metabolism or neurodevelopmental pathways included in heatmaps shown in Figure 5. DAPs identified using default parameters in the DAVID Functional Annotation Tool (28, 29). Pathways with a p-value<0.05 were considered significant and included in analysis. DAPs for energy metabolism and neurodevelopment are included on separate tabs and columns show protein abundances for each sample replicate.
